# Adenomatous polyposis coli (APC) regulates internalization and signaling of the chemorepellent receptor, Roundabout (ROBO) 1

**DOI:** 10.1101/2024.01.12.574717

**Authors:** Yi-Wei Huang, Jonathan St-Germain, Bo Wen Pang, Richard F Collins, Etienne Coyaud, Wenjuan Li, Amir Mohamed, Brian Raught, Sergio Grinstein, Lisa A Robinson

## Abstract

The SLIT-ROBO signaling pathway regulates axon guidance and cell migration, and ROBO1 is a receptor for SLIT ligands. ROBO1 undergoes constitutive endocytosis which is enhanced upon SLIT2 binding, but the molecular mechanisms and functional consequences of this process are not well understood. Using pharmacologic inhibitors and molecular techniques, we found that clathrin-mediated endocytosis is necessary for SLIT2-induced inhibition of cell spreading. To explore the underlying mechanisms, we performed BioID to identify ROBO1-interacting proteins whose association with the cytoplasmic domain of ROBO1 is differentially regulated by SLIT2. We discovered that adenomatous polyposis coli (APC), a multifunctional tumor suppressor, constitutively interacts with ROBO1 but dissociates upon binding of SLIT2 and that this dissociation is necessary for clathrin-mediated endocytosis of ROBO1 and subsequent effects on cell morphology. These findings provide new insights into the functional mechanisms by which SLIT2 binding to ROBO1 effects changes in actin cytoskeletal architecture.

## Introduction

Roundabout (ROBO) receptors are a family of highly conserved axon guidance receptors which together with their specific secreted SLIT ligands play a repulsive role to prevent axons from crossing the midline repeatedly and randomly during neuronal development (Brose et al., 1999) (Kidd et al., 1999). SLIT-ROBO signaling also negatively regulates adhesion, spreading and migration of diverse hematopoietic cell types, including neutrophils, platelets and macrophages (Tole et al., 2009, Patel et al., 2012) and inhibits macropinocytosis in macrophages (Bhosle et al., 2020). SLIT-ROBO regulate actin cytoskeletal remodeling through the inhibition of Rac1 and Cdc42 (Tole et al., 2009) or the activation of Rho A (Bhosle et al., 2020) via the binding of SLIT-ROBO GTPase activating protein (srGAPs) or inactivation of the Rho GTPase-activating protein, myosin IXb (MYO9B), respectively (Wong et al., 2001) (Guerrier et al., 2009).

In the developing nervous system of *Drosophila*, repulsion of midline axons has been demonstrated to require Slit-induced clathrin-mediated endocytosis (CME) of Robo from the cell surface (Chance & Bashaw, 2015) (Howard et al., 2019). In murine cells, endocytosis and recycling of ROBO1 are required for SLIT-induced responses and further sensitization in vertebrate commissural axons upon midline crossing (Kinoshita-Kawada et al., 2019). However, the underlying molecular mechanisms of CME of ROBO in mammalian cells are relatively unexplored. We report here that in mammalian cells SLIT2 binding to ROBO1 induces endocytosis of ROBO1 in a dynamin- and clathrin-dependent manner and that endocytosis of ROBO1 from the cell surface requires the tumor suppressor protein, adenomatous polyposis coli (APC). We found that APC constitutively associates with the intracellular domain of ROBO1, and that binding of SLIT2 to ROBO1 causes APC to dissociate from ROBO1. The functional consequences of APC disengagement from ROBO1 on SLIT2-induced cytoskeletal changes are further discussed.

## Materials and Methods

### Antibodies and reagents

Primary or secondary antibodies were purchased from the following vendors: rabbit anti-human ROBO1 (PA5-29917, Thermo Fisher Scientific, Rockford, IL, USA); rabbit anti-human ROBO1 (HPA052968, Sigma Aldrich, St. Louis, MO); mouse monoclonal anti-human ROBO1 (MAB7118, R&D systems, Minneapolis, MN, USA); mouse monoclonal anti-human APC (CC-1) (ab16794, Abcam Inc, Toronto, ON, Canada); rabbit monoclonal anti-clathrin heavy chain (D3C6) (#4796, Cell Signaling Technology, Danvers, MA, USA); mouse monoclonal anti-clathrin heavy chain (X22) (ab2731, Abcam Inc, Toronto, ON, Canada); mouse monoclonal anti-β-actin (AC004, ABclonal, Woburn, MA, USA); rat monoclonal anti-Flag (637302, Biolegend, San Diego, CA, USA), mouse monoclonal anti-Flag (M2) (F1804, Sigma Aldrich, St. Louis, MO); mouse monoclonal anti-Myc (9B11) (#2276, Cell Signaling, Danvers, MA, USA); Alexa fluor 488-conjugated Fab fragment goat anti-mouse IgG (115-547-003, Jackson ImmunoResearch Laboratories, Inc, West Grove, PA, USA); Alexa fluor 647-conjugated Fab fragment donkey anti-mouse IgG (715-607-003, Jackson ImmunoResearch Laboratories, Inc, West Grove, PA, USA). AF-488-conjugated transferrin was from Molecular Probes by Invitrogen Life Technologies (T13342, Thermo Fisher Scientific, Rockford, IL, USA).

The A Disintegrin and Metalloprotease (ADAM)-10 inhibitor GI 254023X was from Sigma-Aldrich Canada (SML0789, Oakville, ON, CA); the dynamin inhibitor Dyngo4a was from Abcam (ab120689, Abcam Inc, Toronto, ON, CA).

Proximity ligation kits were from Sigma Aldrich (Sigma Aldrich, St.Louis, MO): Duolink® In Situ PLA® Probe Anti-Rabbit PLUS DUO92002; Duolink® In Situ PLA® Probe Anti-Mouse MINUS DUO92004; Duolink® In Situ Detection Reagents FarRed DUO92013.

### DNA plasmid construction

In order to construct pcDNA5 FRT/TO Robo1-WT-BirA*-Flag with restriction sites AscI-NotI, human Robo1 (NP_002932) (Figure s1A) was first sub-cloned from pYFP-C1 ROBO1 WT in pZeoSV2-vector with NheI and HindIII. A linker with NheI-AscI-EcoRV was added in frame to the N-terminus of the Robo1 sequence. A Robo1 C-terminal sequence starting from SalI was substituted by PCR to introduce a NotI restriction enzyme site upstream of HindIII. The Robo1 full length DNA sequence was then sub-cloned into pcDNA5 FRT/TO-BirA*-Flag plasmid with AscI-NotI restriction sites (Figure s1B). An N-terminal Robo1 including the transmembrane domain (M1-R920) was PCR amplified with AscI-NotI sites and sub cloned into pcDNA5 FRT/TO-BirA*-Flag vector to make pcDNA5 FRT/TO Robo1-NTM-BirA*-Flag construct (Figure s1C).

pcDNA3.1zeo+/ cmScarlet-hRobo1 FL was subcloned with NheI-HindIII and mScarlet was PCR cloned with HindIII-Xho1 from pmScarlet C1 and subcloned in frame into pcDNA3.1zeo+ vector Figure s1D). pcDNA3.1zeo+/ nFlag-cmScarlet-hRobo1 was generated by inserting the Flag tag sequence at the N-terminus after the signal peptide sequence of Robo1 with QuikChange II Site-Directed Mutagenesis Kit (Agilent Technologies, Inc, Santa Clara, CA, USA). pcDNA3.1zeo+/ nFlag-intMyc-cmScarlet-hRobo1 was generated by inserting the Myc tag sequence at the extracellular juxta-membrane region of Robo1, downstream of a predicted ADAM10 cleavage site (Q888-Q889), in pcDNA3.1zeo+/ nFlag-cmScarlet-hRobo1 construct (Figure s1 E). The two AP2 binding sites were modified singly or in combination by site-directed mutagenesis in pcDNA3.1zeo+/ cmScarlet-hRobo1 plasmid. Mutation of both AP2 binding sites was established by digesting pcDNA3.1zeo+/ cmScarlet-hRobo1-double AP2 mutant construct with BamH1 and HindIII fragment spanning the two AP2 binding sites and substituting in pcDNA3.1zeo+/ nFlag-intMyc-cmScarlet-hRobo1 WT (Figure s1E and s1F). All DNA plasmid sequences were confirmed by automated sequencing.

The peGFP-C-APC full length DNA plasmid was kindly provided by Dr Angela I. M. Barth (Barth et al., 2002)

All primer sequences for PCR cloning or qPCR are provided in Supplementary Table s1.

### Cell culture

The HEK293A, RAW 264.7 and Cos-7 cell lines were obtained from and authenticated by the American Type Culture Collection (ATCC). Cells were grown in RPMI-1640 Medium containing 5% heat-inactivated fetal calf serum at 37°C with 5% CO2. To generate stable cell lines expressing FL Robo1, pcDNAZeo+/nFlag-intMyc-cmScarlet-Robo1 FL plasmid (Figure s1 E) was transfected in HEK293A cells, cultured, and selected in the presence of zeocin for 2 weeks. Cells were sorted by flow cytometry selecting cells with red fluorescence and a single cell clone (5H9) was obtained by the limited dilution method. HEK293 A cells stably express pcDNAZeo+/nFlag-intMyc-cmScarlet-Robo1 FL construct (Figure s1E) is hereafter referred to as 5H9 cells.

The DNA plasmid nFlag-intMyc-cmScarlet Robo1-AGYA-AAGA (Figure s1 E and s1 F) encoding ROBO1 containing point mutations of both AP2 domains was transfected in HEK293 cells. After cell sorting by flow cytometry and zeocin selection we obtained stable polyclonal cells for subsequent experiments.

Cells stably expressing APC and Robo1 were generated by transfecting peGFP-APC in 5H9 cells, and were selected by culturing in the presence of G418; cell sorting by flow cytometry was performed to select cells demonstrating both green (APC) and red (ROBO1) fluorescence. For comparison, a cell line with GFP-C was established simultaneously.

293 T-REx Flp-In cells stably expressing Robo1-WT-BirA*-Flag or Robo1-NTM-BirA*-Flag were generated using the Flp-In system (Invitrogen, Thermo Fisher Scientific, Rockford, IL, USA) after Hygromycin B selection (Coyaud et al., 2015).

Peripheral blood mononuclear cells (PBMCs) were isolated from the blood of healthy volunteers by density gradient separation using Lympholyte-H (Cedarlane, Burlington, Canada). The cells were cultured in two 10-cm dishes in RPMI 1640 medium supplemented with 10% heat-inactivated FBS, 1% penicillin/ streptomycin/amphotericin B and 10 ng/ml of human M-CSF for 7-12 days (Wong et al., 2016). The protocol for human blood collection from healthy adult donors (#1000070040) was reviewed and approved by The Hospital for Sick Children Research Ethics Board. Informed, written consent was obtained from all participants prior to their participation.

### siRNA gene silencing

siRNA siGENOME targeting human clathrin heavy chain (CLTC) was from Dharmacon (D-004001-02-0005); siRNAs ONTARGETplus human siRNA SMARTpools targeting human APC were purchased from Dharmacon (L-003869-00-0005); siGENOME non-targeting siRNA was from Dharmacon (D-001210-01-20). siRNA transfection was performed using reverse transfection protocols with Lipofectamine RNAiMAX transfection reagent (13778, Life technologies). For CLTC knockdown, cells were transfected with specific siRNA (65 nM) and experiments performed 72 hours later. For APC knockdown, cells were transfected with specific siRNA (50 nM) on Day 0 and Day 2 and experiments performed on Day 4 (96 hours). The siRNA-mediated knockdown efficiency was confirmed by either immunoblotting or qPCR.

### RNA isolation, reverse transcription (RT) and quantitative polymerase chain reaction (qPCR)

Total RNA was isolated from cells using RNeasy® Plus Mini Kit (74034, Qiagen) and RT was performed with 1-2 μg RNA using SuperScript™ IV VILO™ Mastermix (11756050, Thermo Fisher) with the following conditions: 25°C 10 min, 50°C 10 min, 85°C 5 min and products were stored at −20°C until qPCR. qPCR was performed using SYBR™ Green PCR Master Mix kit (4367659, Applied Biosystems). Human GAPDH and green monkey β-actin were used as reference controls. qPCR data were obtained using QuantStudio 3 Real-Time PCR System and analysis was performed using Design and Analysis Software v2 (Applied Biosystems).

All primer sequences for Q-PCR are provided in Table s1.

### Endocytosis assay

Cells were plated in 24-wells with glass coverslips pre-coated with poly-D-lysine (50 μg/mL; MW 70-150 kDa, P6407, Sigma-Aldrich, Canada, Oakville, ON). Cells were treated with the A Disintegrin and Metalloprotease (ADAM)-10 inhibitor, GI-254023X, 10 μM for 16 h to allow ROBO1 to accumulate on the cell surface. Cells were washed twice with PBS to remove the ADAM-10 inhibitor and incubated for 30 min with fresh complete RPMI-1640 medium. Cells were incubated with mouse monoclonal anti-FLAG antibody (1.5 μg/mL) for 1 h at 4°C with slow rotation. The medium was removed and cells were incubated at 37°C with bio-active NSLIT2 (30 nM) or an equimolar concentration of bio-inactive CSLIT2 (Tole et al., 2009, Patel et al., 2012) for different time periods. Endocytosis was arrested by placing plates on ice and cell surface ROBO1 removed using acid wash buffer (100 mM HAC, 0.5 M NaCl in PBS, pH 3.1) for 1 min followed by washing 3 times with PBS. A parallel sample was established for each time point, undergoing only three PBS washes to detect total ROBO1. Cells were fixed with 4% PFA and permeabilized with 0.1% Triton X-100 and 0.1 M glycine for 15 min each. Cells were blocked with 5% goat serum and labeled with AF488 or AF647-Gt-anti mouse IgG fab fragment plus DAPI for 45 min. The percentage of internalized ROBO-1 was calculated based on this equation (Fourgeaud et al., 2003):

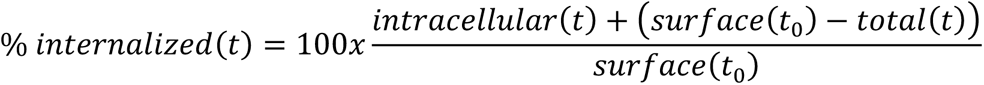

where *intracellular*(*t*) is the fluorescence signal remaining after acid wash at the specified time and represents the endocytosed receptors; *total* (*t*) represents the fluorescence signal after PBS wash including cell surface and internalized signals; *surface*(*t*_0_) − *total*(*t*) represents the endocytosed or degraded receptor fluorescence signal inside cells.

In parallel experiments, 5H9 cells were serum starved for 30 min, placed on ice, and incubated with AF-488 conjugated transferrin (50 μg/mL) for 45 min. Cells were warmed to 37°C and incubated with NSLIT2 or CSLIT2 for different time periods. Endocytosis assays were performed as described above for ROBO-1.

### Cell spreading assay

Human blood monocyte-derived macrophages, RAW264.7 cells or COS-7 cells were grown to 90% confluence, serum starved for 30 min, detached, and exposed to NSlit2 or bio-inactive CSLIT2 or *Δ*D2SLIT2 (Patel et al., 2012, Bhosle et al., 2020) for 15 min. Cells (0.5-1 x 10^5^) were added to poly-D-lysine pre-coated coverslips in 12-well plates for 1 h in serum free RPMI-1640 medium and allowed to attach and spread. Coverslips were fixed with 4% PFA and permeabilized with 0.1 M glycine and 0.1% triton-100. Cells were incubated with AF488-phalloidin or AF488-phalloidin together with DAPI and mounted for immunofluorescence microscopy.

### Proximity ligation assay

COS-7 cells (∼ 3-5 x 10^4^) were grown on 8-well chamber glass slides (BD Falcon 8 chamber tissue culture-treated glass slides, Ref 354118, Corning, NY USA) in serum-free RPMI-1640. Proximity ligation assays (PLA) were performed according to the manufacturer’s instructions. All incubations were performed in a black humidified chamber starting from the blocking step. Briefly, cells were incubated with NSLIT2 or CSLIT2 in serum-free medium for 15 min and samples were washed twice with PBS and fixed with 4% PFA. After washing three times with PBS, cells were permeabilized with 0.1 M glycine, 0.1% TritonX100 in PBS for 15 min, washed again three times, and incubated with the blocking buffer provided by the kit for 1 h at 37°C. Slides were incubated with primary antibodies at 4°C overnight, washed three times with buffer A (0.01 M Tris, 0.15 M NaCl, 0.05% Tween-20, pH 7.4), and incubated with secondary antibody at 37°C for 1 h. Slides were processed for the ligation for 30 min at 37°C, washed twice with buffer A, and amplification was done for 100 min at 37°C. Slides were washed twice with buffer B (0.2 M Tris, 0.1 M NaCl, pH 7.5) and once with 0.01 x wash buffer B. Mounting medium with DAPI (ab104139, abcam) was added to the slides and microscopy performed using a Quorum Spinning Disk Confocal microscope. Imaging data were analyzed by Volocity 6.3 software using the same criteria setting as in Blobfinder software (Allalou and Wahlby, 2009). Interactions were quantified by counting the number of dots per nucleus as well as the intensity of the signal per dot. An increase in intensity is the consequence of a concentration of interactions in the same cellular dots (Gauthier et al., 2015)

### Immunoblotting

Cells were lysed using ice-cold RIPA lysis buffer with protease inhibitor cocktail (Sigma-Aldrich Canada, Oakville, ON). Samples were run on SDS-PAGE, transferred to 0.45 μm PVDF (Millipore) membrane, and probed with specific primary antibodies and the corresponding HRP-conjugated goat anti-mouse or anti-rabbit secondary antibodies. The membrane was washed with TBST, treated with Supersignal^TM^ Chemiluminescent substrate (Thermo Fisher Scientific, Rockford, IL, USA) and visualized on a ChemiDoc MP imaging system (Bio-Rad laboratories, Mississauga, ON, Canada). Band intensity was quantified using ImageJ software, version 1.53k.

### Immunofluorescence microscopy imaging acquisition and analysis

Immunofluorescence microscopy was performed using a Quorum Spinning Disk Confocal microscope Leica DMi8 (Leica microsystems) equipped with a Hamamatsu C9100-13 EMCCD camera and 40× (NA-1.3) oil immersion objectives for endocytosis and cell spreading assays. For proximity ligation assays, a 40× (NA-1.1) water immersion objective was used. For all experiments, 7 to 15 random fields for each experimental condition were imaged as Z-stacked images. Image analyses were performed using Volocity 6.3 software (PerkinElmer, Waltham, MA, USA).

### BioID Sample Processing

BioID (Roux et al., 2012) was performed as described previously (Gingras et al., 2019, Astori et al., 2020). In brief, after selection (DMEM + 10% FBS + 200 μg/mL hygromycin B), two independent replicates of five 150 cm^2^ plates of sub-confluent (60%) BirA*-Flag or ROBO1-BirA*-Flag fusion-expressing cells were incubated for 24 hours in complete medium supplemented with 1 µg/mL tetracycline (Sigma-Aldrich, Canada, Oakville, ON) and 50 µM biotin (BioShop Canada, Burlington, ON). Cells were incubated with purified NSlit2 (5 μg/ml) for 24 hours together with tetracycline. Cells were collected and pelleted (2,000 rpm, 3 minutes), the pellet was washed twice with PBS and dried pellets were snap frozen. The cell pellet was resuspended in 10 mL of lysis buffer (50 mM Tris-HCl pH 7.5, 150 mM NaCl, 1 mM EDTA, 1 mM EGTA, 1% Triton X-100, 0.1% SDS, 1:500 protease inhibitor cocktail (Sigma-Aldrich, Canada, Oakville, ON), 1:1,000 benzonase nuclease (Novagen, Sigma Aldrich, St.Louis, MO) and incubated on an end-over-end rotator at 4°C for 1 h, briefly sonicated to disrupt any visible aggregates, then centrifuged at 45,000 x g for 30 min at 4°C. Supernatant was transferred to a fresh 15 mL conical tube. 30 μL of packed, pre-equilibrated Streptavidin-sepharose beads (GE) were added and the mixture incubated for 3 h at 4°C with end-over-end rotation. Beads were pelleted by centrifugation at 2,000 rpm for 2 min and transferred with 1 mL of lysis buffer to a fresh Eppendorf tube. Beads were washed once with 1 mL lysis buffer and twice with 1 mL of 50 mM ammonium bicarbonate (pH=8.3). Beads were transferred in ammonium bicarbonate to a fresh centrifuge tube and washed two more times with 1 mL ammonium bicarbonate buffer. Tryptic digestion was performed by incubating the beads with 1 µg MS-grade TPCK trypsin (Promega, Madison, WI) dissolved in 200 μL of 50 mM ammonium bicarbonate (pH 8.3) overnight at 37°C. The next morning, 0.5 μg MS-grade TPCK trypsin was added, and beads were incubated 2 additional hours at 37°C. Beads were pelleted by centrifugation at 2,000 x g for 2 min, and the supernatant was transferred to a fresh Eppendorf tube. Beads were washed twice with 150 µL of 50 mM ammonium bicarbonate, and these washes were pooled with the first eluate. The sample was lyophilized and resuspended in buffer A (0.1% formic acid). 1/5th of the sample was analyzed per MS run.

### BioID LC-MS

Peptides were subjected to liquid chromatography (LC) performed on pre-columns (150-mm I.D.) and analytical columns (75-mm I.D.) assembled in-house using fused silica capillary tubing (InnovaQuartz, Phoenix, AZ) packed with 100 Å C18 (Magic, Michrom Bioresources, Auburn, CA). A 120-minute reversed-phase chromatographic gradient (0-100% can, 0.1% HCOOh) running at 250 nL/min on a Proxeon EASY-nLC pump was applied to samples in-line with a hybrid LTQ-Orbitrap Velos mass spectrometer (Thermo Fisher Scientific, Waltham, MA). A parent ion scan was performed in the Orbitrap at a 60,000 resolution (fwhm), with up to the 20 most intense precursors being selected for MS/MS (minimum ion count of 1000 for activation), using standard collision-induced dissociation fragmentation. Dynamic exclusion was activated such that MS/MS of the same m/z (within a range of 15 ppm; exclusion list size = 500) detected twice within 15 seconds were excluded from analysis for 30 seconds. For protein identification, RAW files were converted to the .mzXML format using Proteowizard (Holman et al., 2014), then searched using X!Tandem (Craig and Beavis, 2004) against the human database (Human RefSeq Version 45). Parent ion mass tolerance error of 15ppm and fragment ion mass of 0.4 Da were indicated. No fixed modifications were included, but oxidation (M), acetylation (protein N-terminus) and deamidation (N, Q) were included as potential modifications. Each sample was analyzed using two technical replicates. Data were analyzed using the trans-proteomic pipeline (TPP) (Keller et al., 2005) via the ProHits software suite (Liu et al., 2010). Proteins identified with a Protein Prophet cut-off of 0.9 were analyzed with the SAINT (Choi et al., 2011) algorithm using 4 BirA*-Flag samples as negative controls (12 BirA*-Flag only controls compressed to 4). High confidence interactors were defined as those with bayesian false discovery rate (BFDR) ≤0.01 (table s2). Raw mass spectrometry data have been uploaded to the MassIVE public repository (https://massive.ucsd.edu/) under accession# MSV000092650.

### Statistical analyses

Statistical analyses were performed using GraphPad Prism 8 software (San Diego, CA, USA), and the results are presented as mean ± standard deviation SD). Student’s t-test was used for two-group comparisons, one-way ANOVA was used for multiple-group comparisons with a single independent variable, and two-way ANOVA was used for multiple-group comparisons with two independent variables. Post-hoc multiple comparisons tests (Tukey’s or Sidak’s) were used to compare individual groups. A p-value of less than 0.05 was considered statistically significant.

## Results

### ROBO1 undergoes basal endocytosis which is enhanced by binding of the ligand, SLIT2

It has been reported that SLIT-dependent endocytic trafficking of the Robo receptor is necessary for its repulsive signaling output in *Drosophila* (Chance and Bashaw, 2015). To investigate whether human ROBO1 undergoes endocytosis, a HEK293A cell line stably expressing human ROBO1 fused with an N-terminal FLAG tag, extracellular juxtamembrane Myc tag, and C-terminal mScarlet tag (pcDNAZeo+/nFlag-intMyc-cmScarlet-Robo1) was established, and referred to as 5H9 cells (Figure s1E). The extracellular domain of ROBO1 undergoes constitutive shedding by the metalloprotease, ADAM10 (Coleman et al., 2010). Therefore, to maximize ROBO1 levels on the surface of 5H9 cells, cells were incubated with the ADAM10 inhibitor, GI 254023X, prior to performing endocytosis experiments, and internalization of the FLAG signal was serially monitored (figure 1A, B) (Reiss et al., 2005). For cells exposed to vehicle or to bio-inactive CSLIT2, internalization of ROBO-1 from the cell surface was ≈ 20% at 15 min, 40% at 30 min, and 50% at 60 min (figure 1C). Exposure to bioactive NSLIT2 significantly increased internalization of ROBO1 at all time points, to ≈ 60% at 15 min, 75% at 30 min, and 80% at 60 min (figure 1C; p=0.0002, 0.0001 and 0.0106, respectively, vs vehicle or CSLIT2). To determine whether NSLIT2 enhanced the internalization of cell surface receptors other than ROBO-1, we measured endocytosis of transferrin. At all time points examined, internalization of transferrin was similar for cells exposed to vehicle control, CSLIT2, and NSLIT2 (figure 1 E; p=0.1165, 0.9765 and 0.3331 at 7, 15, 30 min, respectively, vs vehicle, p=0.4012, 0.3585 and 0.2861 vs CSLIT2). Together, these results indicate that exposure of cells to NSLIT2 specifically enhances internalization of ROBO1 from the cell surface but does not stimulate endocytosis globally.

**Figure1.**
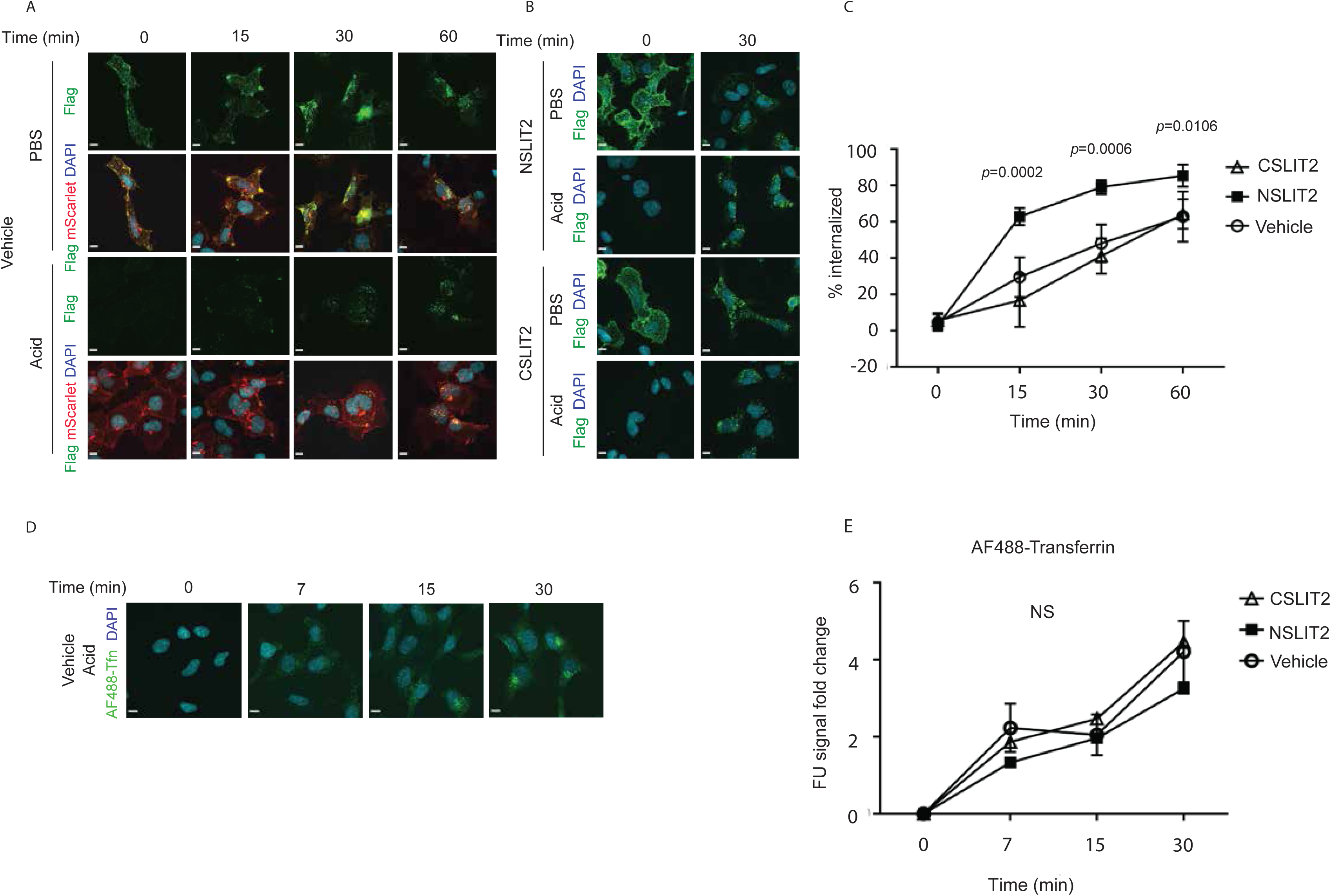
ROBO1 undergoes constitutive endocytosis which is augmented by exposure to NSLIT2. A) 5H9 cells on coverslips were treated with the ADAM10 inhibitor GI 254023X (10 μM) overnight to maximize cell surface levels of ROBO1, then incubated with anti-Flag antibody for 1 h on ice. Cells were warmed to 37 ^ο^C to allow endocytosis. At indicated time points cells were subjected to acid wash or wash with PBS, fixed and permeabilized, then labeled with AF488-conjugated Gt αM secondary antibody. Images were acquired using Quorum Spinning Disk Confocal microscope Leica DMi8. Green, N-terminal Flag-tagged ROBO1; red, C-terminal mScarlet ROBO1. B) Experiments were performed as in (A) but cells were incubated with NSLIT2 (30 nM) or equimolar concentration CSLIT2 prior to initiating endocytosis at 37 ^ο^C. Representative images are shown at 0 min and 30 min time points. C) Experiments were performed as in (B) and the percentage of internalized Flag-tagged ROBO1 was serially measured. Results are from 4 independent experiments. p values were determined using two-way ANOVA. D) Experiments were performed as in (A), incubating cells with AF488-conjugated transferrin to assess endocytosis of transferrin at serial time points. Representative images are shown at indicated time points. E) Experiments were performed as in (D) and the percentage of internalized AF488-Tfn was serially measured. Results are from 3 independent experiments. p values were determined using two-way ANOVA. NS= not significant. Scale bars in each image are 10 μm.

### NSLIT2-induced endocytosis of ROBO1 from the cell surface is clathrin-dependent

In *Drosophila*, Slit-induced endocytosis of Robo and subsequent axonal repulsion have been shown to be dynamin-and clathrin-dependent, and to require AP2-binding motifs present in the cytoplasmic domain of ROBO-1 (Chance and Bashaw, 2015). We, therefore, used Dyngo-4a, a potent dynamin inhibitor (McCluskey et al., 2013, Basagiannis et al., 2021) to investigate the role of dynamin in mediating endocytosis of ROBO-1 in mammalian cells. As expected, inhibition of dynamin attenuated internalization of transferrin (Figure s2A, 2B). Dynamin inhibition decreased endocytosis of ROBO-1 by 55% (figure 2A, 2B; p=0.0049 for Dyngo-4a 30 μM vs vehicle; p=0.006 for Dyngo-4a 40 μM).

**Figure 2.**
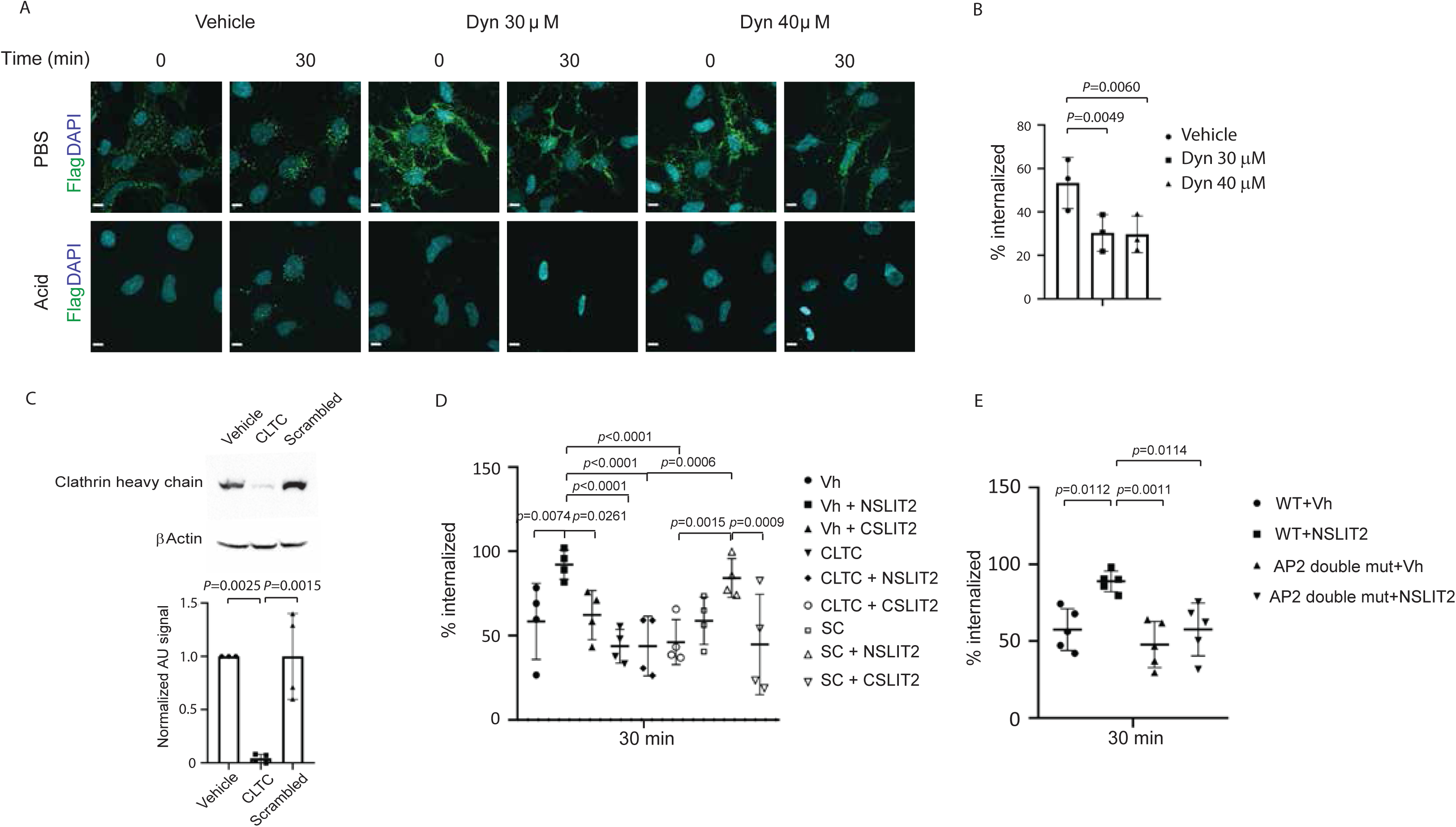
NSLIT2-induced endocytosis of ROBO1 is clathrin dependent. A) 5H9 cells were incubated with the dynamin inhibitor, Dyngo-4a (Dyn), at the concentrations indicated and endocytosis of ROBO1 assessed as in figure 1A. Representative images are shown at the indicated time points. B) Experiments were performed as in (A) and the percentage of ROBO internalized at 0 min and 30 min were measured. Results were from three independent experiments. p values were determined using two-way ANOVA. C) Cells were incubated with specific siRNA targeting clathrin heavy chain (CLTC) or with scrambled, non-targeting siRNA. After 4 days, cell lysates were obtained and Immunoblotting performed to assess levels of clathrin. Representative image and quantification of three independent experiments. D) Silencing of clathrin was performed as in (C) and endocytosis of ROBO1 assessed as in figure 1B. Results were from 4 independent experiments. E) An allele of Robo1 in which both AP2 binding motifs were mutated was expressed in cells and endocytosis experiments performed as in (C). Results from 5 independent experiments. Scale bars 10 μm.

To examine the role of clathrin in endocytosis of ROBO-1 in mammalian cells, we used siRNA to knock down clathrin to less than 10% of the levels in vehicle-treated cells or in cells exposed to scrambled siRNA (figure 2C; p=0.0025 and 0.0015, respectively) and confirmed that knockdown of clathrin prevented endocytosis of transferrin (Figure s2C, 2D; p=0.0003 clathrin vs. vehicle, p=0.0117 clathrin vs scrambled). In clathrin-deficient cells, NSLIT2-induced endocytosis of ROBO-1 was markedly lower than in cells exposed to control vehicle or to scrambled siRNA (figure 2D; p<0.0001 vs vehicle; p= 0.0006 vs scrambled siRNA).

Similar to Robo in *Drosophila* (Chance and Bashaw, 2015), we found that human ROBO-1 also contains two AP2-binding motifs (Yxxφ) (Collins, 2002) at the C terminus between the CC2 and CC3 domains of the protein (1311 YGYI 1314 and 1413 YAGL 1416). To examine the role of these AP2-binding motifs in endocytosis of mammalian ROBO-1, we established HEK293A cells stably expressing mutant alleles of both AP2-binding sites (1311AGYA1314 and 1413AAGA1416). Exposure of cells expressing wild-type ROBO-1 to NSLIT2 enhanced endocytosis by 36% (figure 2E, p=0.0112), whereas this effect was abolished in cells expressing the mutant alleles of AP2 (figure 2E, p=0.682, WT vs AP2 mutant; p=0.676 WT+NSLIT2 vs. AP2 mut+NSLIT2). These results demonstrate that in mammalian cells, as in *Drosophila* (Chance and Bashaw, 2015), SLIT2-enhanced endocytosis of ROBO-1 is dependent on dynamin, clathrin, and AP2-binding motifs in the intracellular domain of ROBO-1.

### Inhibition of cell spreading by NSLIT2 requires clathrin-mediated endocytosis of ROBO-1

We and others have previously shown that NSLIT2 inhibits spreading of diverse cell types, including neutrophils, platelets, and human and murine macrophages (Patel et al., 2012, Bhosle et al., 2020, Tole et al., 2009). We first confirmed that in primary human monocyte-derived macrophages and the murine RAW264.7 macrophage cell line, exposure of cells to NSLIT2 significantly decreased cell spreading, and that this response was not observed following exposure of cells to bio-inactive CSLIT2 or ΔD2SLIT2, which lacks the ROBO-1-binding domain (Bhosle et al., 2020) (Figure s3A-D; human macrophages: p<0.0001 vs CSLIT2, p=0.0004 vs ΔD2SLIT2; RAW264.7: p=0.0065 vs CSLIT2, p=0.012 vs ΔD2SLIT2). We similarly observed that NSLIT2 significantly inhibited cell spreading of COS-7 cells (figure 3A, 3B, p=0.0498 vs CSLIT2, p=0.0272 vs ΔD2SLIT2). To investigate the role of dynamin in NSLIT2-induced inhibition of cell spreading, we incubated COS-7 cells with the dynamin inhibitor, Dyngo-4a, and observed that NSLIT2 was no longer able to inhibit cell spreading (figure 3C, 3D; p=0.0437). In cells expressing either or both alleles of mutant AP2, NSLIT2 failed to inhibit cell spreading, and the effect was most pronounced for cells expressing both mutant alleles (figure 3E-3G; p=0.0212 vs AGYA, p=0.0053vs AAGA, p<0.0001 vs double mutant). Taken together, these results suggest that in mammalian cells, NSLIT2-induced effects on cell morphology require clathrin-mediated endocytosis.

**Figure 3.**
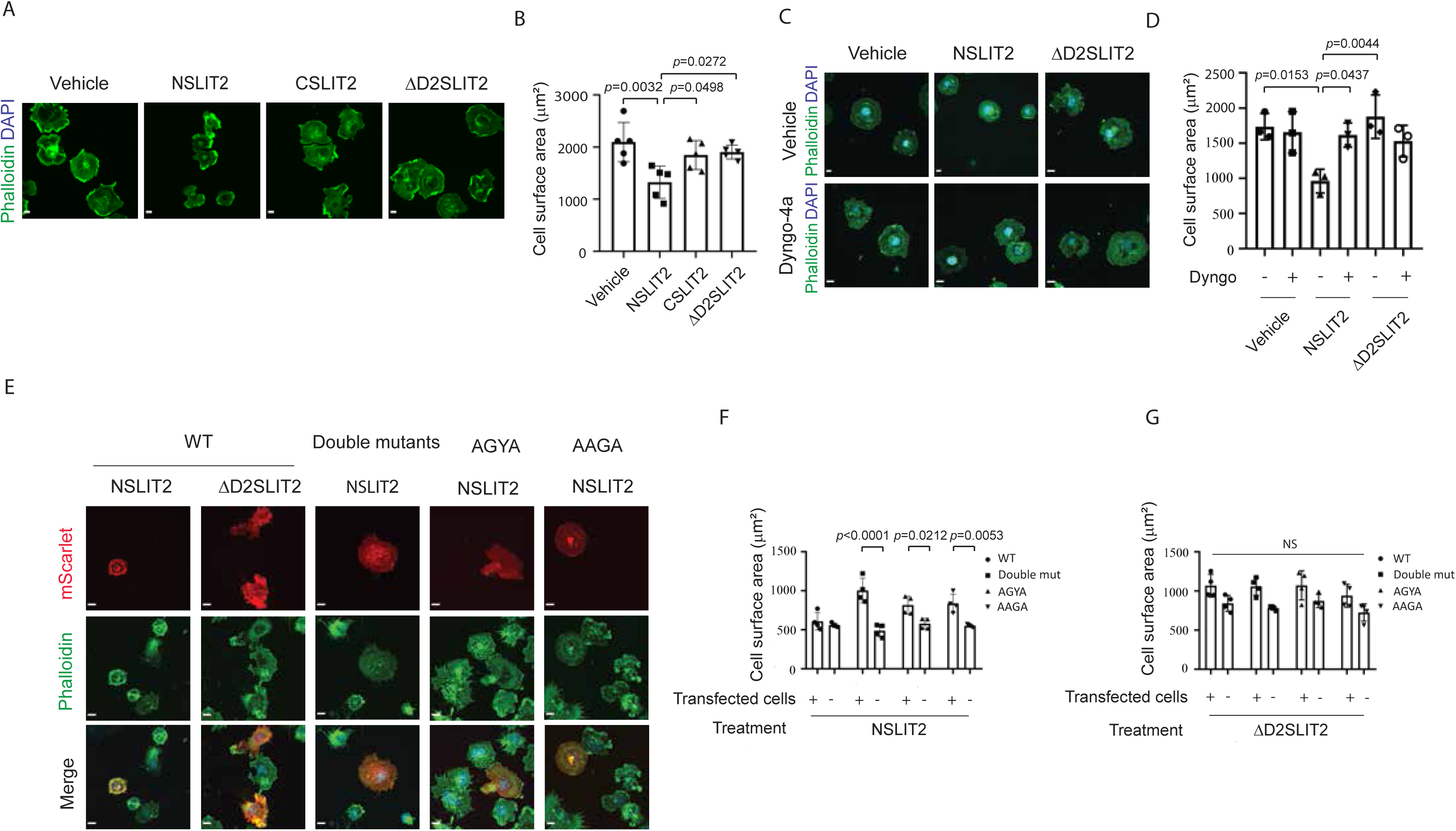
Clathrin-mediated endocytosis of ROBO1 is required for NSLIT2-induced inhibition of cell spreading. A) COS-7 cells were incubated with NSLIT2, CSLIT2, or ΔD2SLIT2 for 15 min at 37 °C, added onto poly-D-lysine-coated coverslips and allowed to spread for 1 h. Cells were fixed, permeabilized, and incubated with AF-488-conjugated phalloidin (green). B) Experiments were performed as in (A) and cell surface area was measured. Results from 5 independent experiments. C) Cells were pre-incubated with dynamin inhibitor, Dyngo-4a (25 μM), for 30 min and experiments performed as in (A). Representative images. D) Experiments were performed as in (C) and cell surface area was measured. Results from 4 independent experiments. Results among groups were compared using one-way ANOVA. E) Cells were transfected with plasmids encoding C-mScarlet ROBO1 FL or ROBO1 containing mutations of AP2-binding regions. After 24 h, cell spreading assays were performed as in (A), incubating cells with NSLIT2 or ΔD2SLIT2. Representative images shown. Scale bar 10 μm. F) Spreading of cells treated with NSLIT2 was measured. G) Spreading of cells treated with ΔD2SLIT2 was measured. Results from 4 Independent experiments. Results among groups were compared with two-way ANOVA.

### Exposure of cells to NSLIT2 differentially regulates protein interactions with ROBO1

To elucidate the mechanism by which NSLIT2 affects cells morphology through clathrin-mediated endocytosis, we sought proteins differentially associated with ROBO1 by performing proximity-dependent biotin identification (BioID) expressing the full-length cBirA ROBO1 protein in HEK 293 Flp-In cells, in the presence and absence of NSLIT2. Cells expressing a form of ROBO1 lacking the cytoplasmic C-terminal region (cBirA ROBO1 NTM) (Figure s1C) or the BirA-Flag tag alone were used as controls.

Through the BioID analysis, we identified a number of proteins which strongly associated with full-length ROBO1 (ROBO1 FL) but not with ROBO1 lacking the intracellular domain (ROBO NTM) (Table 1). These proteins included srGAP2, a known effector of SLIT-ROBO signaling (Charrier, 2012) (Wong et al., 2001) (Guerrier et al., 2009), and two of the five components of the WAVE complex, namely, NCKAP1 (NAP1) and CYFIP1 (Sra), previously shown to interact with ROBO1 to modulate cytoskeletal remodeling (Table 1) (Chen et al., 2014) (Chaudhari, 2021). Interrogation of the BioID data also revealed around 30 candidate proteins which associated with ROBO1 FL under basal conditions but no longer associated with ROBO1 FL after cell exposure to NSLIT2 (Table s2). None of the candidate proteins interacted with the form of ROBO1 lacking the intracellular domain (ROBO NTM; Table s2). These data indicate that candidate proteins identified associate with the intracellular domain of ROBO1 under basal conditions, and that exposure to NSLIT2 reverses this association. This list of candidate proteins includes NCKAP1 and CYFIP1, as well as other proteins associated with actin cytoskeletal remodeling, such as DLG5, MAP4K4, PHACTR4, RDX, PAK4, FARP1, MARK3 and NCK1 (Table 1) (Matsumine et al., 1996) (Liu et al., 2014) (Vitorino et al., 2015) (Gao et al., 2016) (Allen et al., 2004) (Morales et al., 2004) (Won et al., 2019) (Cheadle and Biederer, 2012) (Sandi et al., 2017) (Li et al., 2001) (Yang and Bashaw, 2006). Unexpectedly, we detected a significant association between the intracellular domain of ROBO1 and APC, and observed that this interaction disappeared after exposure of cells to NSLIT2 (Table 1).

**Table 1.** Exposure to NSLIT2 differentially regulates protein interactions with ROBO1. Shown is a short-list of statistically significant (BFDR≤0.01) proximity interacting-proteins of full-length, wild-type ROBO1 identified from the BioID performed in T-REx Flp-In HEK293 cells. Values for each protein across different samples represent total spectral matches of assigned peptides. The Baysian false discovery rate (BFDR) was calculated by comparing the spectral counts of each protein in the bait samples with their counts in the negative control samples (BirA*flag). HEK293 Flp-In cells expressing cBirA ROBO1 were incubated with NSLIT2 or vehicle control for 24 hours together with teracycline and the results were compared to samples from cells expressing a form of ROBO1 lacking the cytoplasmic C-terminal region (cBirA ROBO1 NTM).

| Gene Symbc | Gene name | ROBO1_WT-BirA*-Flag |  |  |  |  |  |  |  | ROBO1_NTM-BirA*-Flag |  |  |  |  |  |  |  |
| --- | --- | --- | --- | --- | --- | --- | --- | --- | --- | --- | --- | --- | --- | --- | --- | --- | --- |
|  |  | (-) NSLIT2 |  |  |  | (+) NSLIT2 |  |  |  | (-) NSLIT2 |  |  |  | (+) NSLIT2 |  |  |  |
|  |  | Rep1 | Rep2 | SUM | BFDR | Rep1 | Rep2 | SUM | BFDR | Rep1 | Rep2 | SUM | BFDR | Rep1 | Rep2 | SUM | BFDR |
| ROBO1 | *roundabout guidance receptor 1* | 1806 | 1877 | 3683 | NA | 1164 | 1153 | 2317 | NA | 479 | 488 | 967 | NA | 506 | 511 | 1017 | NA |
| BirA | *E. coli biotin ligase* | 1813 | 2130 | 3943 | NA | 977 | 964 | 1941 | NA | 452 | 446 | 898 | NA | 451 | 480 | 931 | NA |
| DLG5 | *discs, large homolog 5 (Drosophila)* | 27 | 24 | 51 | ≤0.01 | 0 | 0 | 0 | >0.01 | 0 | 0 | 0 | >0.01 | 0 | 0 | 0 | >0.01 |
| CYFIP1 | *cytoplasmic FMR1 interacting protein 1 (Sra)* | 19 | 16 | 35 | ≤0.01 | 0 | 0 | 0 | >0.01 | 0 | 0 | 0 | >0.01 | 0 | 0 | 0 | >0.01 |
| APC | *adenomatous polyposis coli* | 13 | 18 | 31 | ≤0.01 | 0 | 0 | 0 | >0.01 | 0 | 0 | 0 | >0.01 | 0 | 0 | 0 | >0.01 |
| MAP4K4 | *mitogen-activated protein kinase kinase kinase kinase 4* | 10 | 15 | 25 | ≤0.01 | 0 | 0 | 0 | >0.01 | 0 | 0 | 0 | >0.01 | 0 | 0 | 0 | >0.01 |
| PHACTR4 | *phosphatase and actin regulator 4* | 10 | 6 | 16 | ≤0.01 | 0 | 0 | 0 | >0.01 | 0 | 0 | 0 | >0.01 | 0 | 0 | 0 | >0.01 |
| RDX | *radixin* | 7 | 6 | 13 | ≤0.01 | 0 | 0 | 0 | >0.01 | 0 | 0 | 0 | >0.01 | 0 | 0 | 0 | >0.01 |
| NCKAP1 | *NCK-associated protein 1 (NAP1)* | 17 | 21 | 38 | ≤0.01 | 0 | 4 | 4 | >0.01 | 0 | 0 | 0 | >0.01 | 0 | 0 | 0 | >0.01 |
| PAK4 | *p21 protein (Cdc42/Rac)-activated kinase 4* | 4 | 3 | 7 | ≤0.01 | 0 | 0 | 0 | >0.01 | 0 | 0 | 0 | >0.01 | 0 | 0 | 0 | >0.01 |
| MARK3 | *MAP/microtubule affinity-regulating kinase 3* | 3 | 2 | 5 | ≤0.01 | 0 | 0 | 0 | >0.01 | 0 | 0 | 0 | >0.01 | 0 | 0 | 0 | >0.01 |
| FARP1 | *FERM, RhoGEF (ARHGEF) and pleckstrin domain protein 1 (chondrocyte-derived)* | 13 | 11 | 24 | ≤0.01 | 4 | 3 | 7 | >0.01 | 0 | 0 | 0 | >0.01 | 0 | 0 | 0 | >0.01 |
| SRGAP2 | *SLIT-ROBO Rho GTPase activating protein 2* | 15 | 13 | 28 | ≤0.01 | 5 | 4 | 9 | >0.01 | 0 | 0 | 0 | >0.01 | 0 | 0 | 0 | >0.01 |

As expected, the proteins associated with ROBO1 lacking the cytoplasmic C-terminus displayed little overlap with the full-length ROBO1, yielding only proximity interactions with plasma membrane, ER and vacuolar membrane proteins not observed in the full-length ROBO1 interactome.

Together, these results provide a human ROBO1 BioID interactome. Consistent with increased intracellular trafficking, this interactome changes in response to exposure to NSLIT2.

### ROBO1 interacts with adenomatous polyposis coli (APC) and exposure to NSLIT2 disrupts this interaction

Validation of this interaction by co-immunoprecipitation of endogenous proteins was not possible due to a lack of suitable antibodies specifically for IP. Instead, we used Proximity ligation assay (PLA), a highly sensitive and powerful tool to assess the interactions of two proteins less than 40 nm apart (Weibrecht et al., 2010) (Soderberg et al., 2006) (Schweitzer et al., 2000) (Gauthier et al., 2015). To validate the assay, we first verified a strong association between ROBO-1 labeled with mouse and rabbit antibodies directed to human ROBO-1 (figure 4A, 4B). We next detected significant association between ROBO-1 and clathrin, in keeping with previous reports that ROBO-1 undergoes CME (figure 4E, 4F, p<0.0001). We also detected significant association between ROBO-1 and APC in unstimulated cells (figure 4A, 4B, p<0.0001). Importantly, this association was disrupted by exposure of cells to NSLIT2, but not to bio-inactive CSLIT2 (figure 4C,4D; p<0.0001 NSLIT2 vs. vehicle; p<0.0001 NSLIT2 vs CSLIT2). Together, these results suggest that ROBO1, APC and clathrin may form a complex and this complex may be present in clathrin-coated pits. Exposure to NSLIT2 leads to dissociation of APC from ROBO1, potentially allowing it to undergo clathrin-mediated endocytosis.

**Figure 4.**
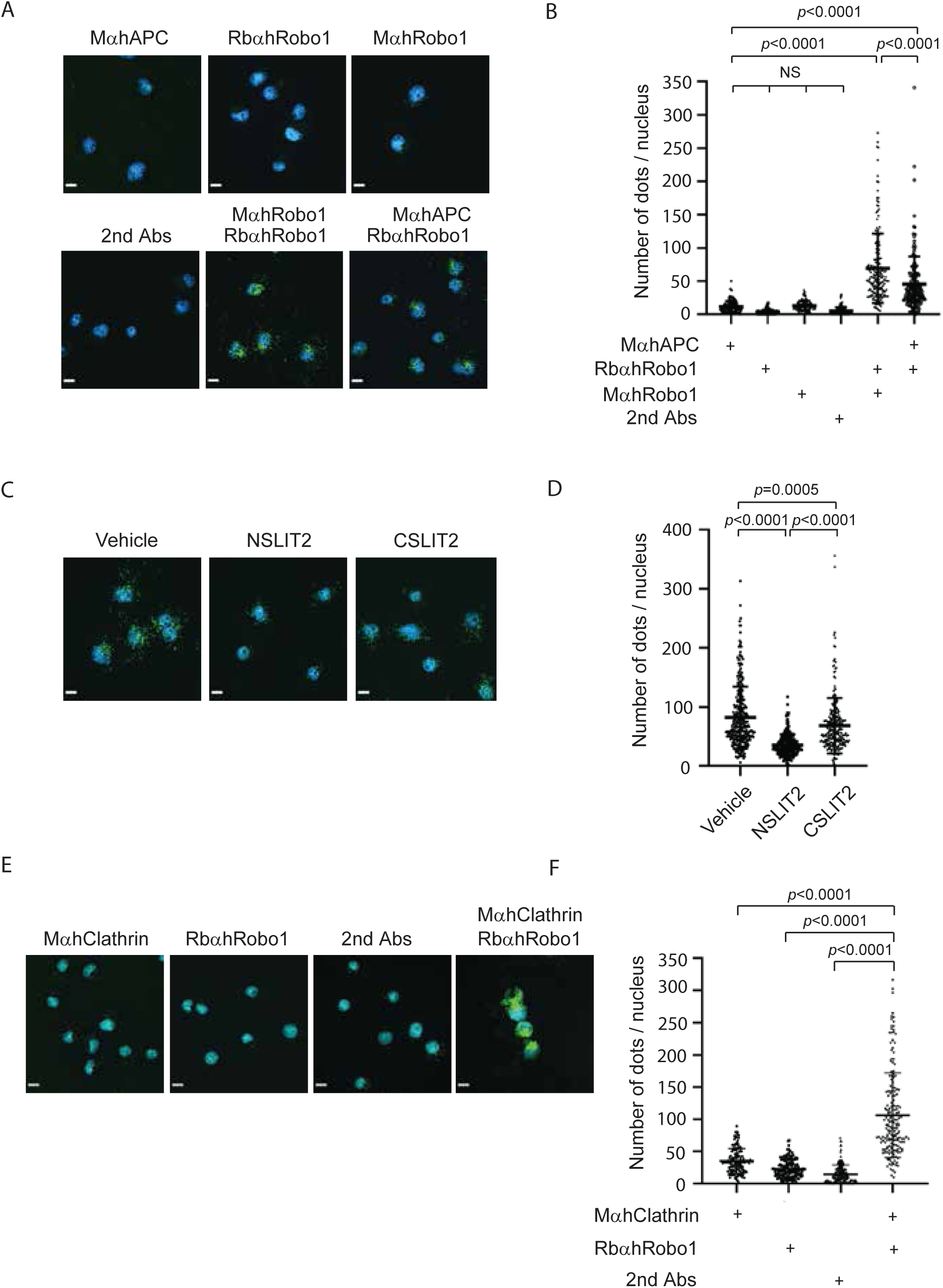
ROBO1 interacts with adenomatous polyposis coli (APC) and clathrin. A) To detect interactions between ROBO1 and APC, proximity ligation assays were performed incubating COS-7 cells with mouse anti-ROBO1 antibody recognizing the N-terminal region of Robo1 (MAB7118, R&D systems), rabbit anti-ROBO1 antibody recognizing the C-terminal region of ROBO1 (HPA052968, Sigma Aldrich), mouse anti-APC antibody, and corresponding secondary antibodies. Representative images are shown. B) Experiments were performed as in (A) and the number of dots per nucleus determined. Results from 3 independent experiments. C) Cell were incubated with NSLIT2 or CSLIT2 and proximity ligation assays performed as in (A). Representative images. D) Experiments were performed as in (C) and the number of dots per nucleus determined. E) To detect interactions between ROBO1 and clathrin, proximity ligation assays were performed using antibodies directed to ROBO1 (HPA052968, Sigma Aldrich) and clathrin (X22, ab2731, Abcam Inc). Representative images are shown. F) Experiments were performed as in (E) and the number of dots per nucleus determined. Results from 3 independent experiments analyzed using one-way ANOVA. Scale bars in images are 10 μm.

### APC modulates NSLIT2-induced endocytosis of ROBO1

To investigate the role of APC in the endocytosis of ROBO-1, we used siRNA to knock down APC in 5H9 cells to less than 10% (figure 5A). In cells incubated with non-targeting scrambled siRNA, exposure to NSLIT2 resulted in significantly more internalization of ROBO-1 than in cells incubated with vehicle or with bio-inactive CSLIT2 (figure 5B). In cells deficient in APC, the basal level of endocytosis was greater than for cells expressing APC (figure 5B; p=0.0078), and endocytosis was similar in the presence and absence of NSLIT2 (figure 5B; p=0.7863). In cells over-expressing APC by ≈30-fold (figure 5C; p=0.0001), the basal level of endocytosis of ROBO-1 was similar to that of cells expressing normal levels of ROBO-1 (figure 5D; p=0.7539). Exposure of such APC over-expressing cells to NSLIT2 failed to stimulate endocytosis, in contrast to the NSLIT2-enhanced endocytosis seen in cells with normal levels of APC (figure 5D; p=0.0031). Together, these results suggest that APC is an endogenous inhibitor of endocytosis for ROBO-1 at the cell surface.

**Figure 5.**
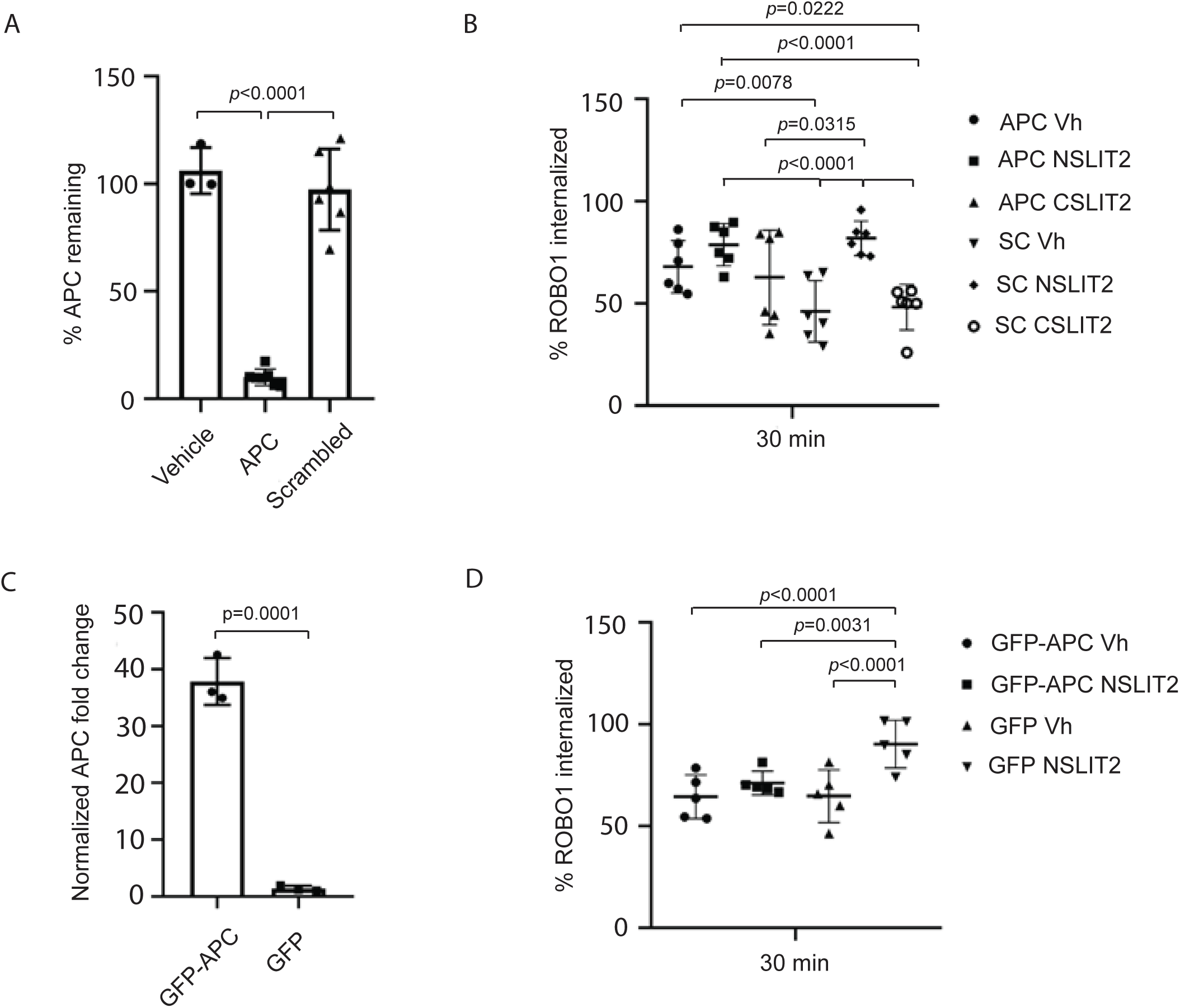
APC modulates NSLIT2-induced endocytosis of ROBO1. A) 5H9 cells were transfected with siRNA specifically targeting APC or with scrambled siRNA. After 4 days, RNA was extracted and APC expression measured using qPCR. Results from 3 to 5 independent experiments were analyzed using one-way ANOVA. B) APC was silenced as in (A) and endocytosis assays performed as in figure 1B. Results from 6 independent experiments were analyzed using two-way ANOVA and Sidak’s multiple comparisons test. C) 5H9 cells were stably transfected to express GFP-tagged APC (GFP-APC) or GFP alone. RNA was extracted and APC expression measured using qPCR. Results from 3 independent experiments analyzed using Students t-test. D) ROBO1 endocytosis assay was done using the cells as shown in (C). The plot was generated from 5 independent experiments. Experiments were analyzed using Two-way ANOVA and Sidak’s multiple comparisons test.

### APC is necessary for NSLIT2-induced inhibition of cell spreading

Having observed that APC inhibits NSLIT2-induced endocytosis of ROBO1, we questioned whether APC is required for the inhibitory effects of NSLIT2 on cell spreading. In COS-7 cells, as in 5H9 cells, we found that a targeted siRNA reduced levels of APC to 25-30% of the levels detected in vehicle-treated cells or in cells exposed to non-targeting scrambled siRNA (figure 6A). In cells with reduced APC, NSLIT2 was no longer able to inhibit cell spreading compared to vehicle-treated cells and cells exposed to scrambled siRNA (figure 6B and 6C; p<0.0001). Taken together, these results suggest that APC is an endogenous inhibitor of endocytosis of a distinct pool of cell surface ROBO1 that is sensitive to the actions of NSLIT2, and that APC is required for NSLIT2-induced inhibition of cell spreading.

**Figure 6.**
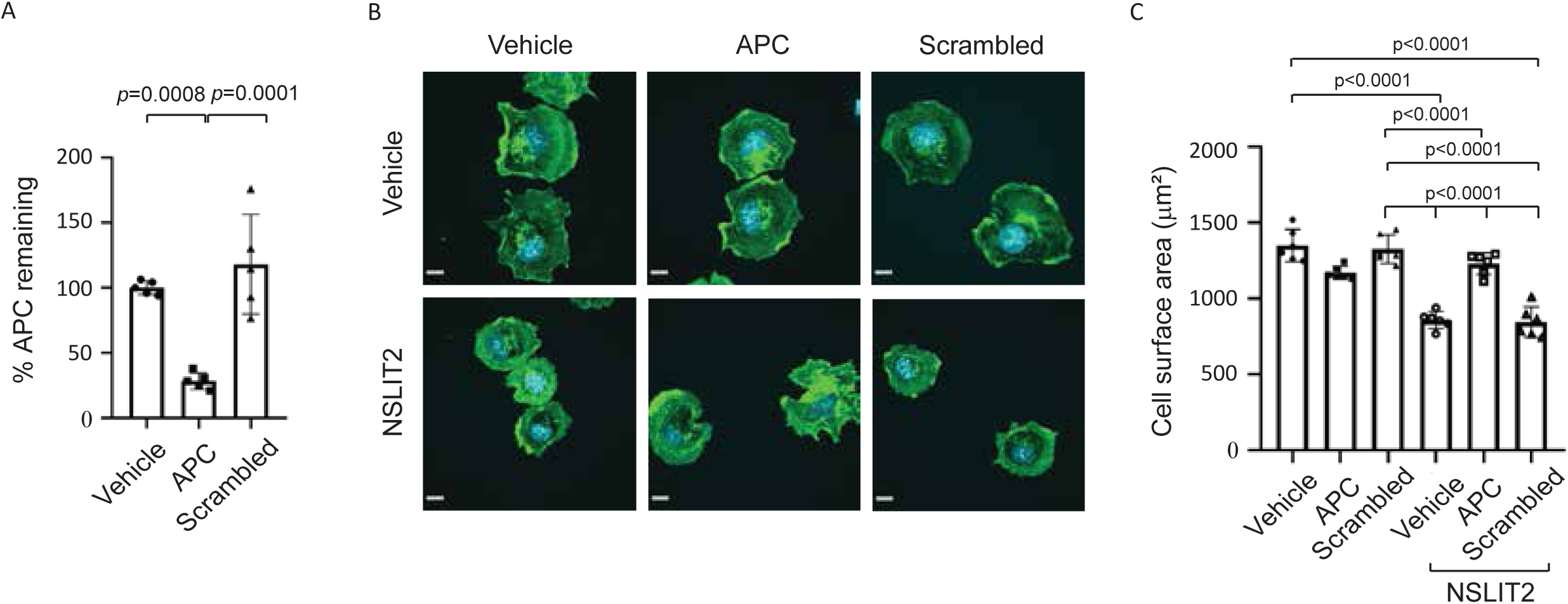
APC is required for NSLIT2-induced inhibition of cell spreading. A) COS-7 cells were transfected with siRNA targeting APC or with scrambled siRNA. After4 days, RNA was extracted and APC expression assessed using Q-PCR. Results from 5 independent experiments were analyzed using one-way ANOVA. B) COS-7 cells were transfected with siRNA targeting APC or with scrambled siRNA. After 4 days, cell spreading assays were performed as in in figure 3A. Representative images. Scale bars are 10 μm C) Experiments were performed as in (B). Results from 5 independent experiments analyzed using one-way ANOVA.

## Discussion

Endocytosis is an essential process that allows cells to regulate the levels of receptors on their surface, control the signaling pathways activated by extracellular ligands, and internalize essential nutrients (Sorkin and von Zastrow, 2009) (McMahon and Boucrot, 2011). ROBO receptors also undergo endocytosis. It has been reported in *Drosophila* that robo1 endocytosis is clathrin-dependent and is essential for receptor activation and proper midline axon repulsion by allowing recruitment of the downstream effector, Son of Sevenless (Chance and Bashaw, 2015). In mammalian cells, it has similarly been reported that endocytosis and recycling of ROBO1 are required for SLIT-induced responses regulating midline crossing of commissural axons (Kinoshita-Kawada et al., 2019). Our data demonstrate that in a variety of mammalian cells ROBO1 also undergoes SLIT2-induced endocytosis that is dynamin- and clathrin-dependent, and that CME is necessary for NSLIT2-induced inhibition of cell spreading. To further identify ROBO1-interacting proteins in an unbiased, sensitive, and specific manner we used BioID, enabling us detection of both physiologically relevant and weak protein-protein interactions (Roux et al., 2012) (Roux et al., 2018). Using BioID, we identified around 11 candidate proteins that interact with the cytoplasmic region of ROBO1 under basal conditions and that dissociate from ROBO1 upon exposure of cells to NSLIT2 (Table 1). ROBO1-interacting proteins identified include important actin cytoskeletal regulatory proteins, among them two of the five components of the WAVE complex, namely NCKAP1 (NAP1) and CYFIP1 (Sra) (Takenawa and Suetsugu, 2007) (Chen et al., 2014). Unexpectedly, we noted that one of the strongest “hits” identified was APC, originally identified as an important tumor suppressor in the human colon and which is conserved across species (Aoki and Taketo, 2007).

APC is a multi-domain protein that contains binding sites for numerous proteins, including microtubules, the Wnt/Wg pathway components β-catenin and axin, the cytoskeletal regulators EB1 and IQGAP1, and the Rac guanine-nucleotide-exchange factor (GEF) Asef1. Activation of the canonical Wnt signal plays an essential role in tumorigenesis caused by *APC* mutations. APC plays additional roles in other cellular processes, such as cell migration, adhesion, and chromosome segregation (Kroboth et al., 2007) (Aoki and Taketo, 2007) (Nelson and Nathke, 2013) (Hedman et al., 2015) (Watanabe et al., 2004) (Kawasaki et al., 2000).

Using proximity ligation assays, we observed that Robo1, APC, and clathrin interact with one another, suggesting that they may form a complex in clathrin-coated pits. This is in keeping with studies demonstrating that APC interacts with the AP2-clathrin complex (Saito-Diaz et al., 2018). We further noted that knock down of APC resulted in a trend to more constitutive endocytosis compared to that seen in cells exposed to scrambled siRNA, suggesting that APC negatively regulates endocytosis. When cellular levels of APC were intact, exposure of cells to NSLIT2 resulted in significantly enhanced endocytosis, as well as reduced cell spreading, but these NSLIT2-induced effects were lost in cells lacking APC. These data are in keeping with reports that under basal conditions APC exists in complex with clathrin to prevent constitutive, ligand-independent, clathrin-dependent signalosome formation and Wnt pathway activation (Saito-Diaz et al., 2018). We further observed that cells expressing overabundant amounts of APC undergo similar rates of endocytosis compared to cells expressing usual amounts of APC, but that unlike in cells expressing normal levels of APC, exposure to NSLIT2 failed to enhance endocytosis in cells over-expressing APC. Together, these results suggest that a limited pool of APC associates with clathrin complexes to constitutively inhibit endocytosis, and that NSLIT2 causes disengagement of this pool of APC from ROBO1, thereby promoting increased internalization of ROBO1. These results are in keeping with crystallographic structural data demonstrating that binding of NSLIT2 to ROBO1 does not promote changes in oligomerization of ROBO1, suggesting instead that binding of NSLIT2 induces a conformational change in ROBO1 that enables subsequent transduction of intracellular signals (Barak et al., 2019). We found that even in the absence of NSLIT2, ROBO1 undergoes some degree of constitutive endocytosis. It has been reported that TGF-β receptors undergo internalization through both clathrin coated pits and caveolae-lipid raft internalization pathways. Clathrin-dependent internalization into the early endosome promotes the activation of the TGF-β signalling cascade, whereas the caveolin-1-positive lipid raft compartment is required for receptor degradation (Di Guglielmo et al., 2003). Other authors reported that nerve growth factor (NGF) leads to the redirection of a pool of p75 neurotrophin receptor (p75NTR) into clathrin-coated pits and internalization of p75NTR via clathrin-mediated endocytosis. However, exposure to NGF does not enhance the internalization or degradation of p75NTR, which undergoes a rapid dynamin-dependent and clathrin-independent recycling process in motor neurons (Deinhardt et al., 2007). Similarly, at low doses, epidermal growth factor (EGF) has been shown to direct its receptor, EGFR, to a clathrin-mediated pathway of internalization resulting in activation of downstream targets. However, at high concentrations EGF promotes EGFR uptake via a clathrin-independent mechanism, resulting in increased receptor degradation (Sigismund et al., 2005)

In keeping with observations from other groups, our findings suggest that ROBO1 may undergoes clathrin-dependent and clathrin-independent endocytosis. A separate pool of ROBO1 is acutely responsive to NSLIT2, suggesting that in the basal state, this pool of ROBO1 may exist in complex with APC, AP2 and clathrin, inhibiting endocytosis of ROBO1. Following binding of NSLIT2, ROBO1 may undergo conformational changes (Barak et al., 2019) that actively disengage APC from the ROBO1 intracellular domain. In turn, this process may dissociate APC from known interacting regulatory proteins, such as the Rac-specific GEFs, Asef1 and Asef2, and/or IQGAP, which inactivate Rac1 and Cdc42, subsequently inhibiting actin cytoskeletal re-arrangements needed for cell spreading and polarized migration (Watanabe et al., 2004) (Hedman et al., 2015). The clathrin-independent endocytosis pathway may distinctly participate in the recycling or degradation of ROBO1. Further studies are needed to determine how internalization of ROBO1 through these two distinct pathways is differentially regulated, and how NSLIT2 selectively affects a distinct pool of ROBO1.

## Author contributions

Y.W.H. designed and performed experiments, analyzed and interpreted the data, and prepared the manuscript.

J.S.G. analyzed and interpreted the data

B.W.P. performed experiments and analyzed data

R.C. performed experiments and analyzed data

E.C. designed and performed experiments, analyzed and interpreted the data

W.L. analyzed and interpreted the data

A.M. performed experiments and analyzed data

B.R. interpreted the data and critically reviewed the manuscript

S.G. and L.A.R. designed experiments, interpreted data, and prepared the manuscript.

## Abbreviations

AGM: African green monkey
AP2: AP2 adaptor complex
APC: adenomatous polyposis coli
BioID: proximity-dependent biotin identification
CLTC gene: Clathrin heavy chain
CME: clathrin-mediated endocytosis
CYFIP1: cytoplasmic FMR1 interacting protein 1 (Sra)
FARP1: FERM, ARH/RhoGEF and pleckstrin domain protein 1
MARK3: MAP/microtubule affinity-regulating kinase 3
MAP4K4: mitogen-activated protein kinase kinase kinase kinase 4
NCK1: NCK adaptor protein 1
NCKAP1: NCK-associated protein 1 (NAP1)
PAK4: p21 protein (Cdc42/Rac)-activated kinase 4
PHACTR4: phosphatase and actin regulator 4
RDX: radixin
SC: Scrambled
WT: wild type

## Figure legends

**Figure s1.**
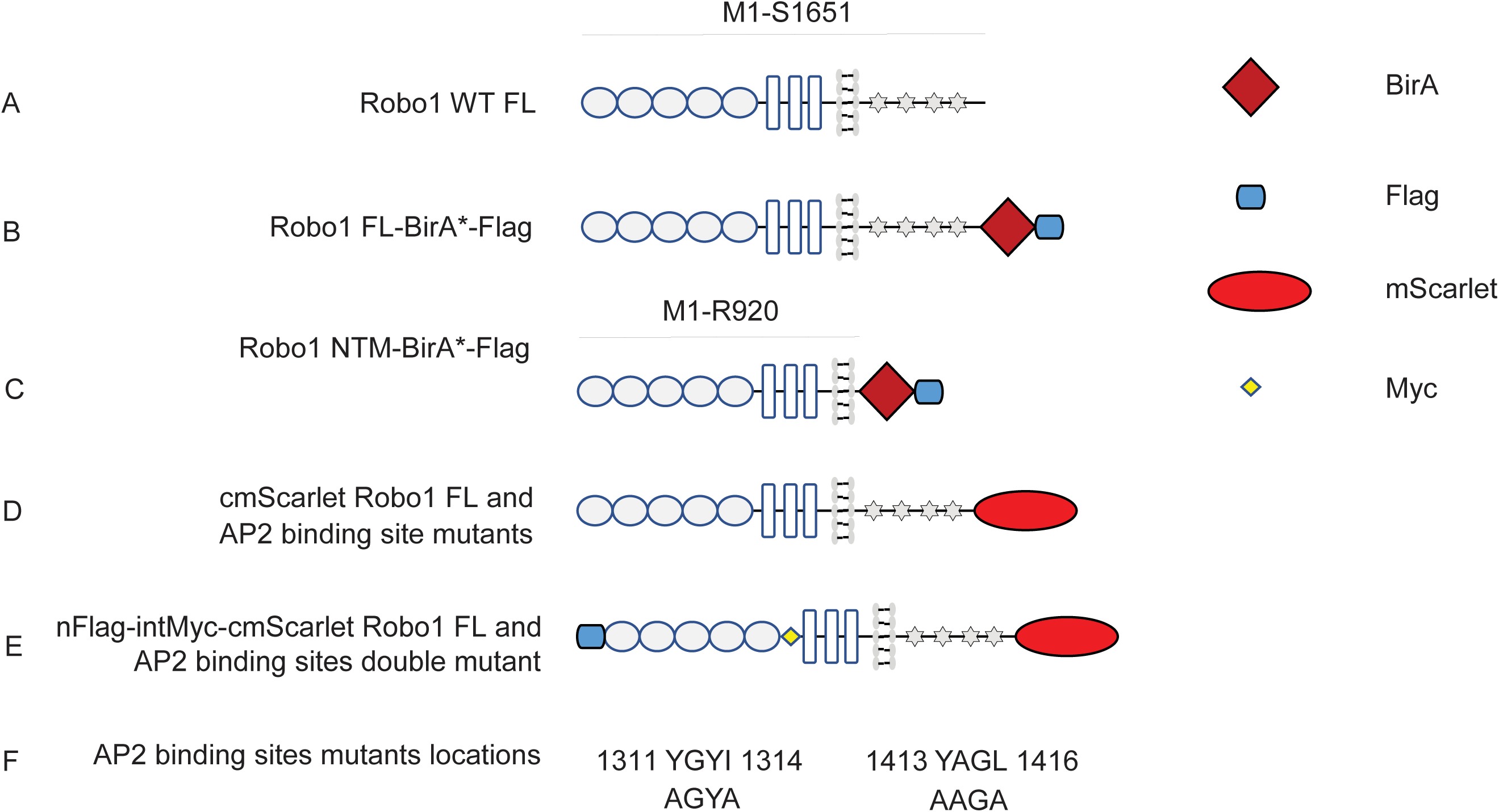
Schematic of Robo1 cDNA plasmids. A) represents the original YFP-tagged full-lenth ROBO1 plasmid (pYFP C1-ROBO1 FL) and an intermediate state of pzeo-/ ROBO1 FL used to generate plasmids for BioID experiments. B) represents the pcDNA5 FRT/TO cBirA-Flag ROBO1 FL plasmid. C) represents pcDNA5 FRT/TO cBirA-Flag ROBO1 NTM (N-terminal mutant) plasmid. D) represents pcDNA zeo+/cmScarlet ROBO1 FL, pcDNA zeo+/cmScarlet ROBO1 AGYA or pcDNA zeo+/cmScarlet ROBO1 AAGA, or pcDNA zeo+/cmScarlet ROBO1 AP2 double mutant plasmids. E) represents pcDNA zeo+/nFlag-intMyc-cmScarlet ROBO1 FL or pcDNA zeo+/nFlag-intMyc-cmScarlet ROBO1 AP2 double mutant plasmids. F) Location of AP2-binding motifs in ROBO1.

**Figure s2.**
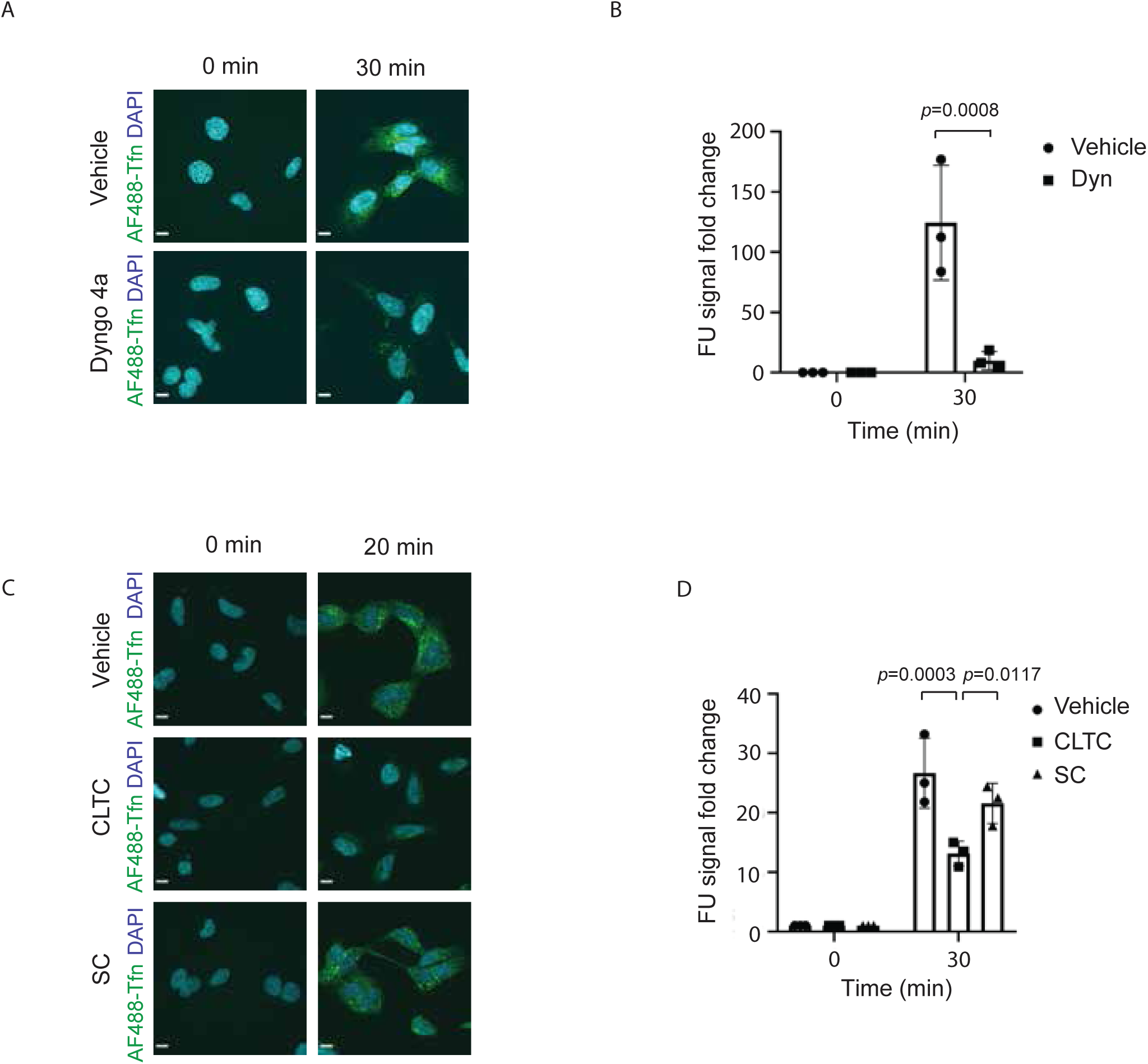
Endocytosis of transferrin is dynamin- and clathrin-dependent. A) 5H9 cells were incubated with the dynamin inhibitor, Dyngo-4a, and endocytosis of AF488-transferrin assessed as in figure 1D. Representative images. B) Experiments were performed as in (A). Results from 3 independent experiments were analyzed using Two-way ANOVA and Sidak’s multiple comparisons test. C) Experiments were performed as in figure 2C. Representative images. D) Experiments were performed as in figure 2D. Results from 3 independent experiments were analyzed using Two-way ANOVA and Sidak’s multiple comparisons test. Scale bars in images are 10 μm.

**Figure s3.**
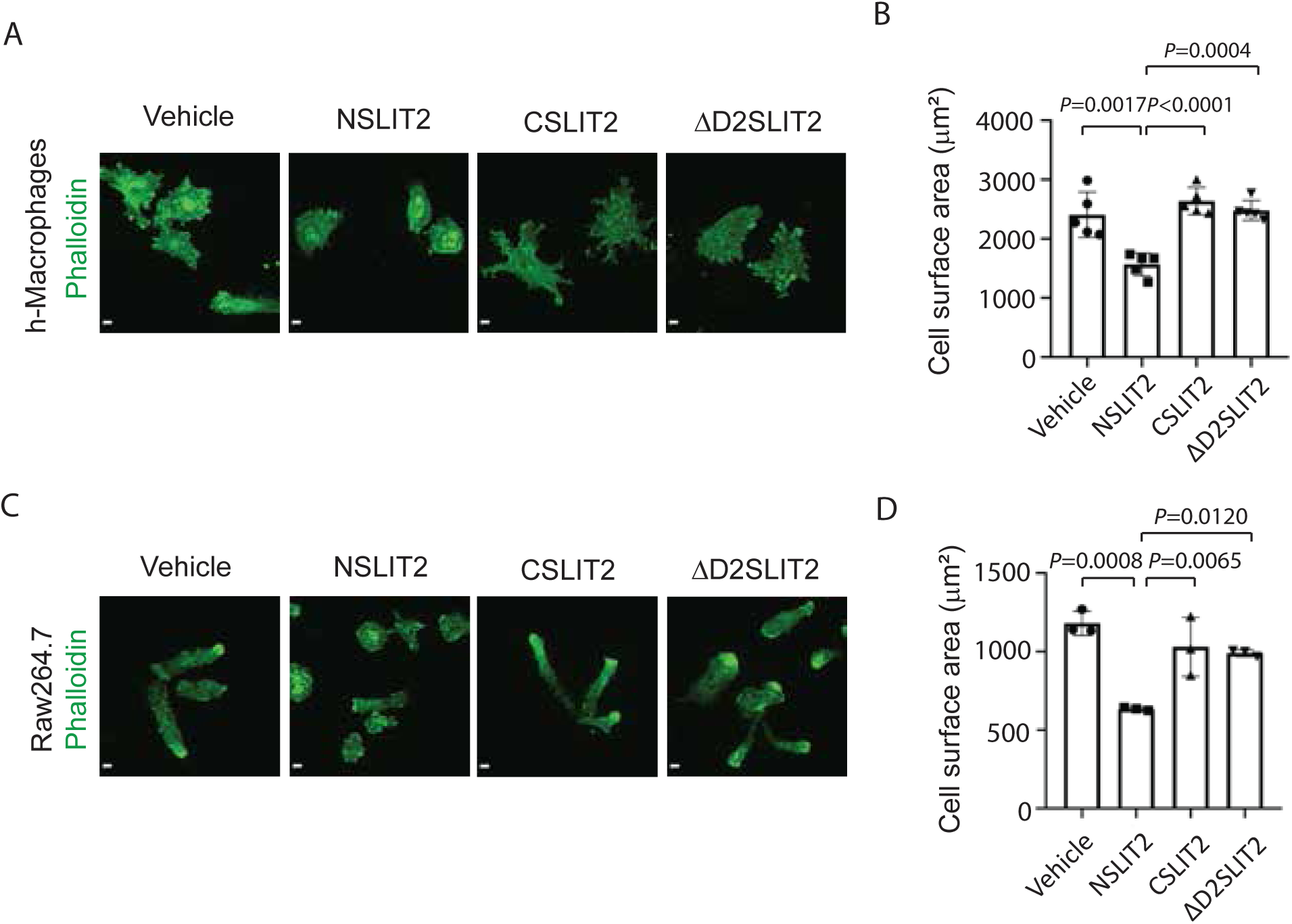
NSLIT2 inhibits spreading of human monocyte-derived macrophages and RAW 264.7 cells. A) Experiments were performed as in figure 3A using primary human monocyte-derived macrophages. Representative images. B) Experiments were performed as in figure 3B. Results from 5 independent experiments analyzed using one-way ANOVA. C) Experiments were performed as in figure 3A using RAW 264.7 cells. Representative images. D) Experiments were performed as in figure 3B. Results from 3 independent experiments were analyzed using one-way ANOVA. Scale bars in images are 10 μm.

---

**Table s1.**
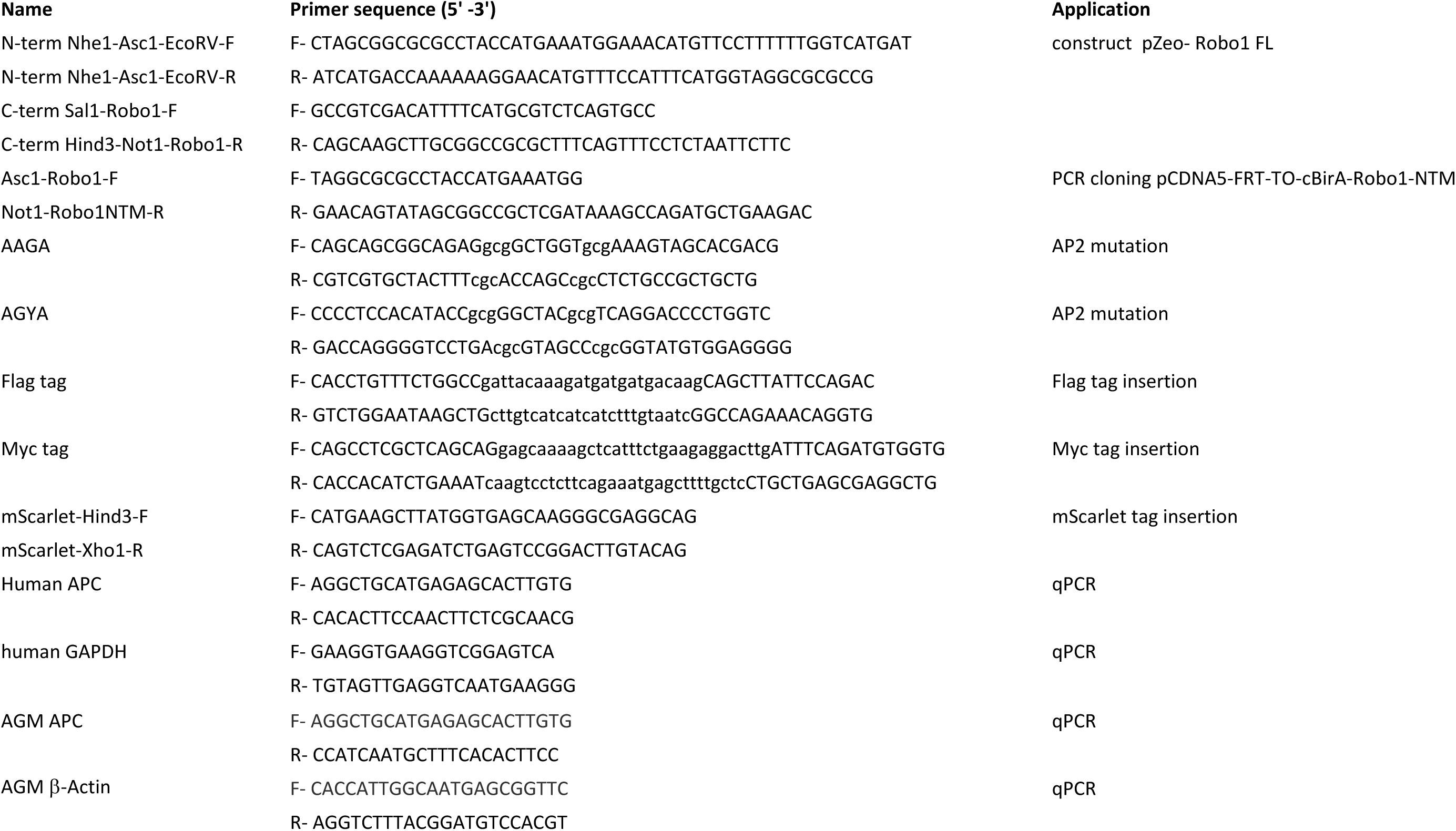
Primers used for cDNA plasmid construction and qPCR experiments.

**Table s2.**
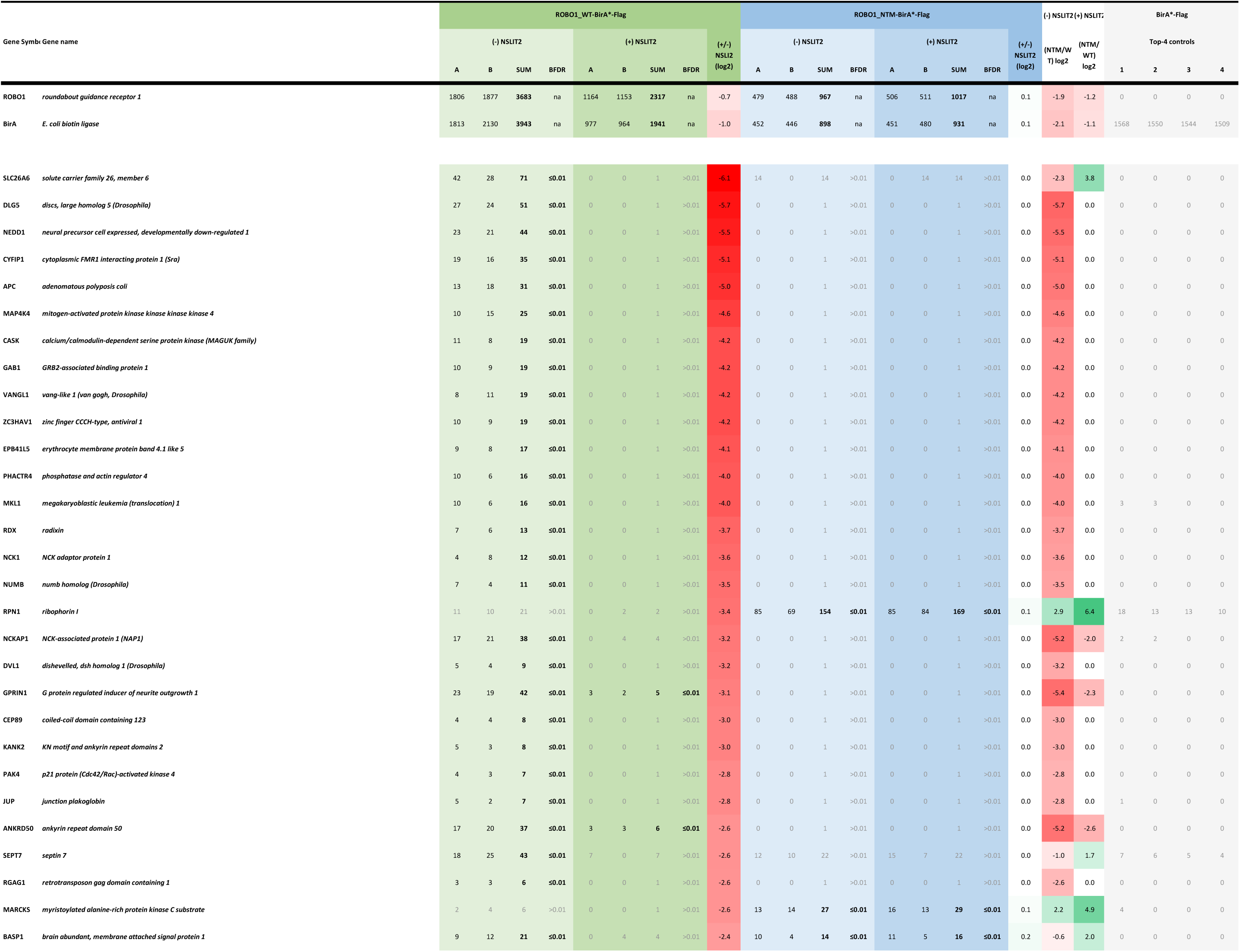
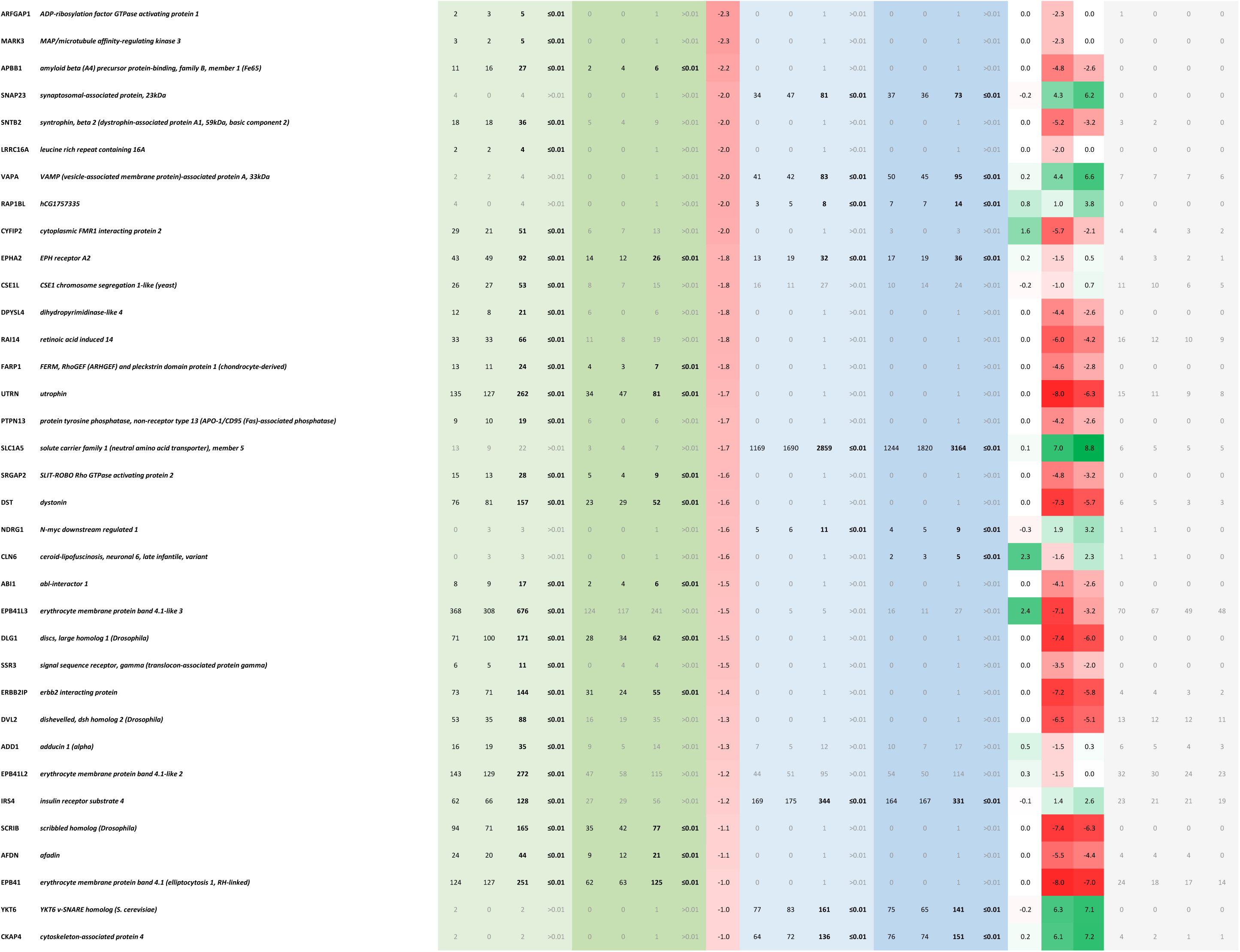
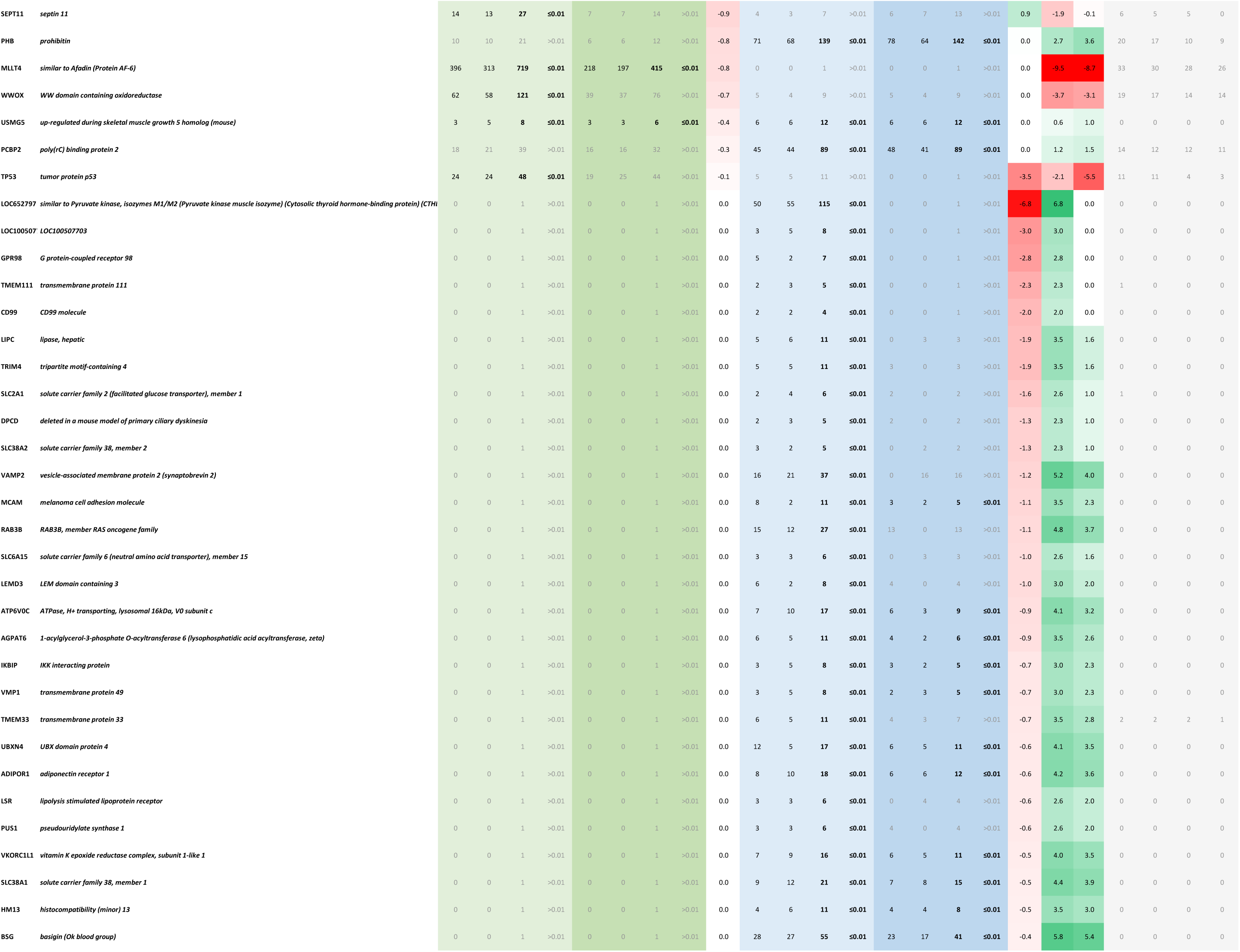
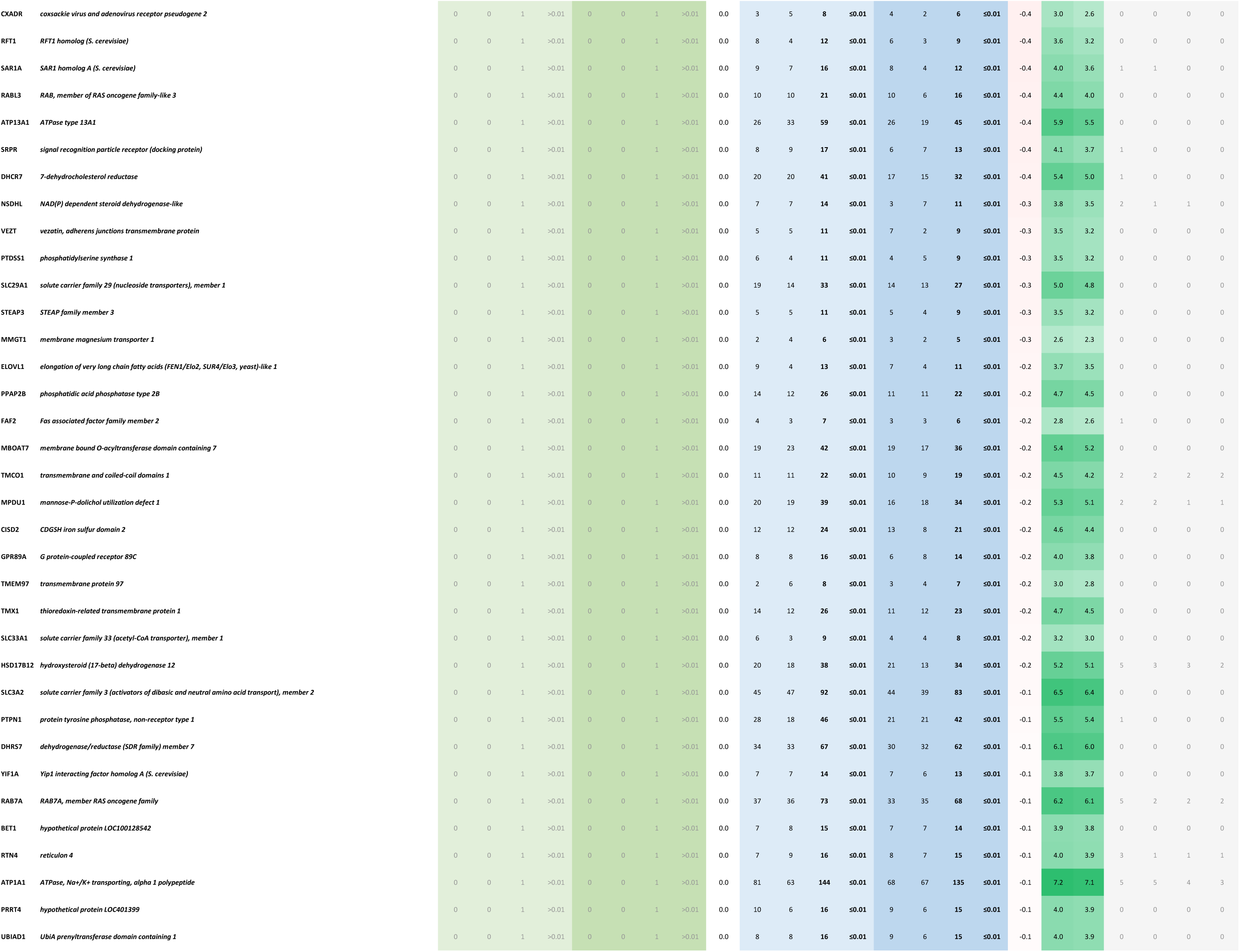
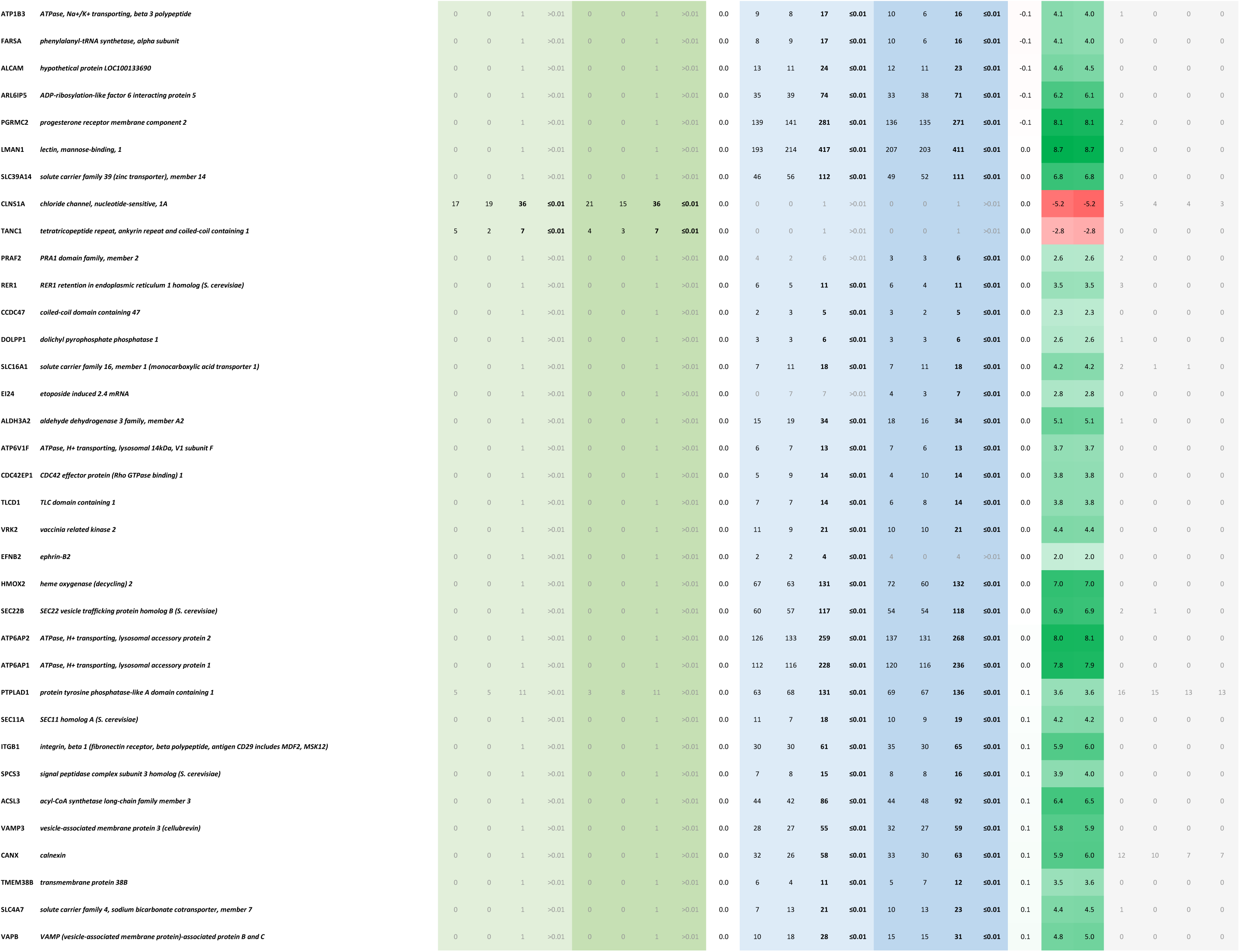
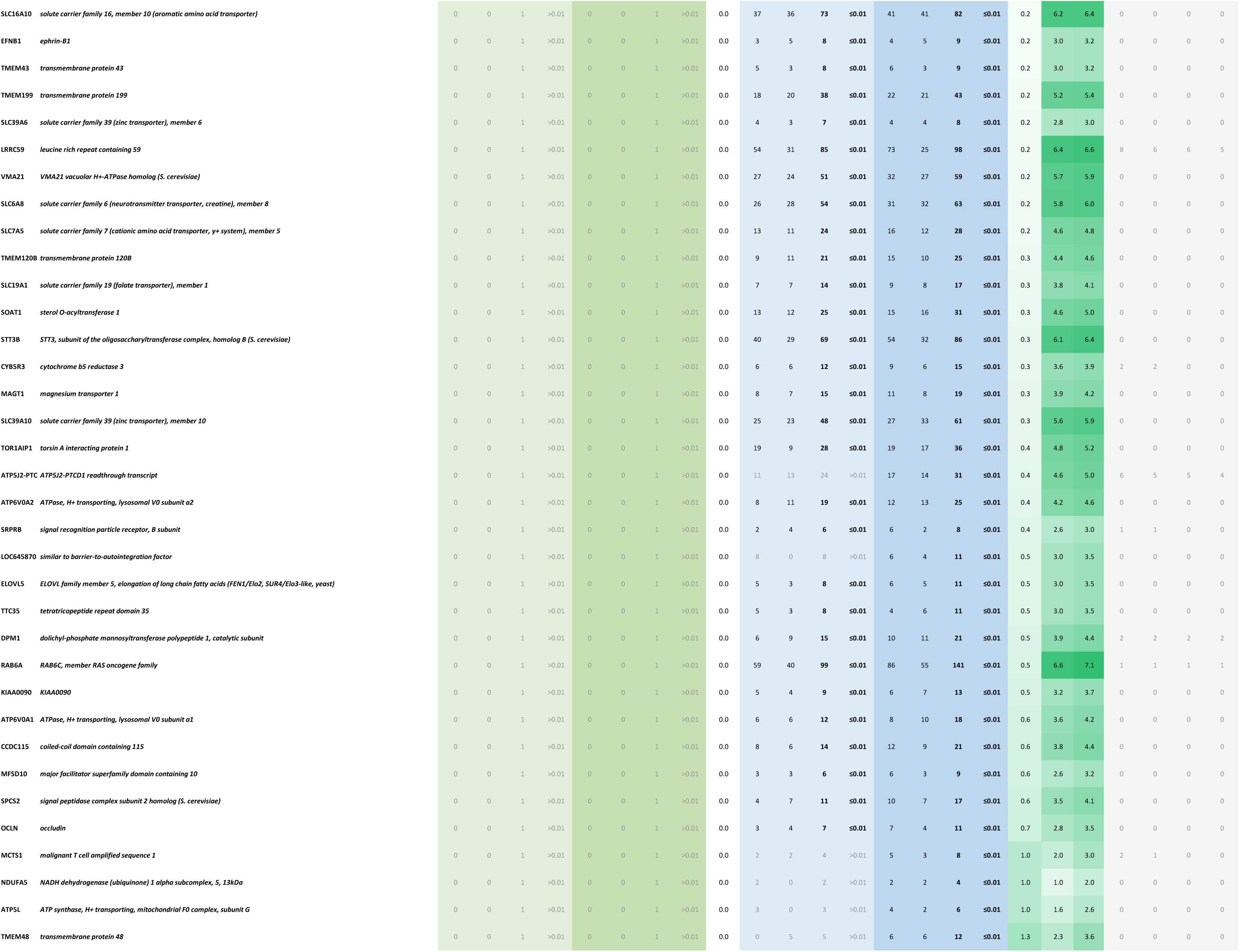
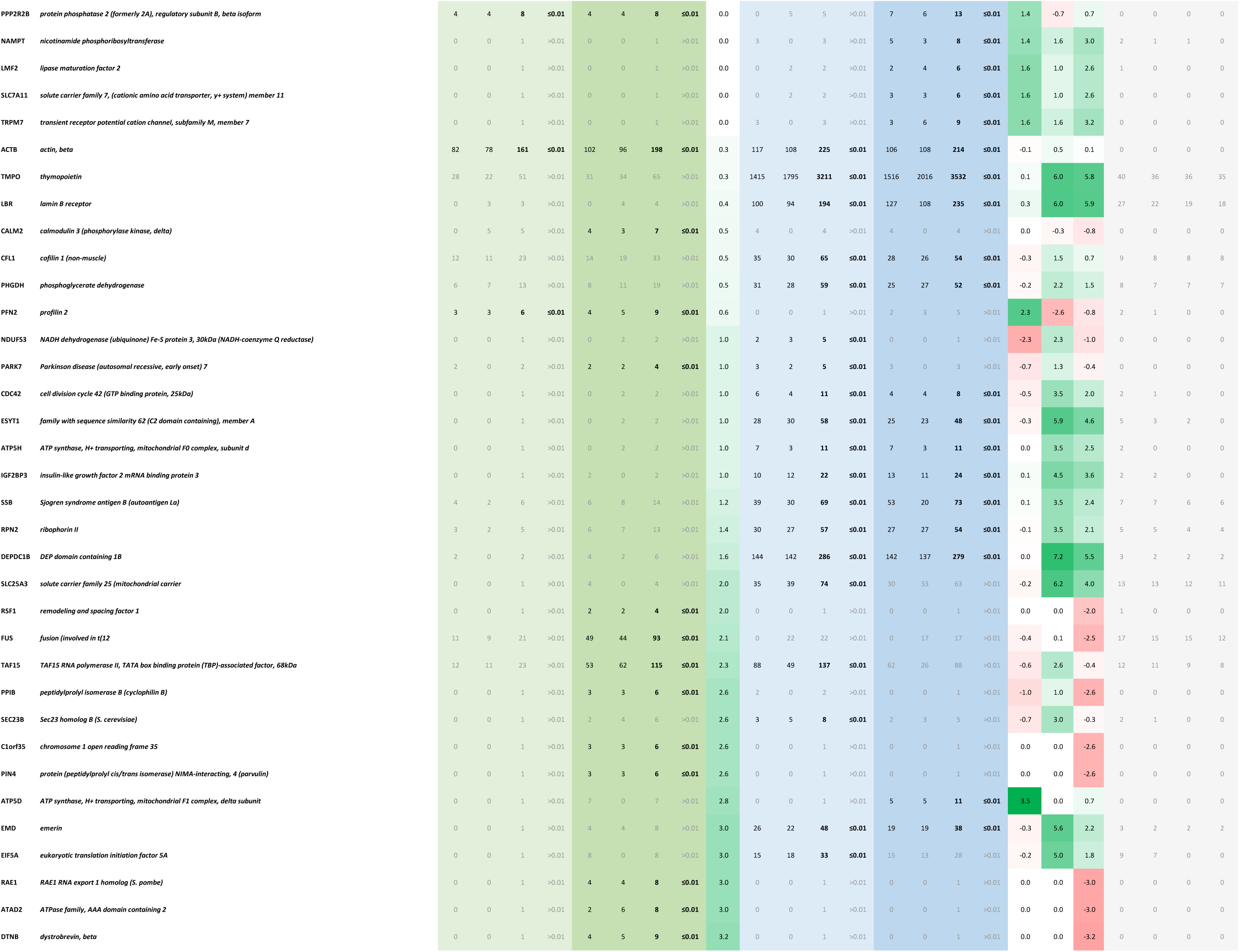
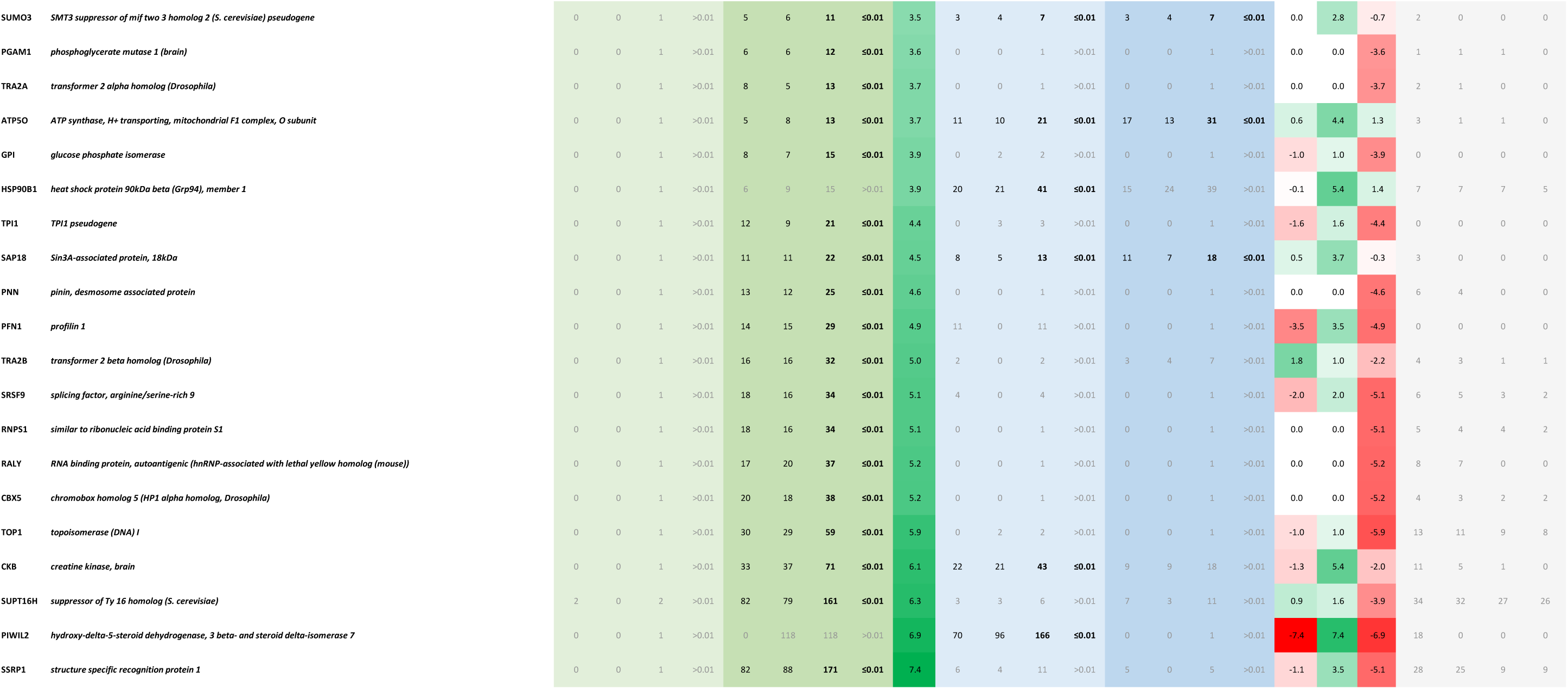
ROBO1 BioID. List of all proteins identified from the proximity-dependent biotin labeling experiment performed in T-REx Flp-In HEK293 cells with indicated fusion proteins as baits. Values for each protein across different samples represent total spectral matches of assigned peptides. The Baysian false discovery rate (BFDR) was calculated by comparing the spectral counts of each protein in the bait samples with their counts in the negative control samples (BirA*flag). The respective fold changes are expressed as the log2 ratio of the summed spectral counts from the replicates.

## Acknowledgments

We would like to thank Paul Paroutis and Kimberly Lau from the SickKids imaging facility for their assistance in imaging acquisition and analysis. This work was supported by grants from Canadian Institutes of Health Research (PJT-169167, MOP-111083, and MOP-136896) and Natural Sciences and Engineering Research Council of Canada (NSERC RGPRN 2017-06460) to L.A.R. L.A.R. holds a Canadian Research Chair in Vascular inflammation and Kidney Injury.

